# Structure-function coupling and decoupling during movie-watching and resting-state: Novel insights bridging EEG and structural imaging

**DOI:** 10.1101/2024.04.05.588337

**Authors:** Venkatesh Subramani, Giulia Lioi, Karim Jerbi, Nicolas Farrugia

## Abstract

The intricate structural and functional architecture of the brain enables a wide range of cognitive processes ranging from perception and action to higher-order abstract thinking. Despite important progress, the relationship between the brain’s structural and functional properties is not yet fully established. In particular, the way the brain’s anatomy shapes its electrophysiological dynamics remains elusive. The electroencephalography (EEG) activity recorded during naturalistic tasks is thought to exhibit patterns of coupling with the underlying brain structure that vary as a function of behavior. Yet these patterns have not yet been sufficiently quantified. We address this gap by jointly examining individual Diffusion-Weighted Imaging (DWI) scans and continuous EEG recorded during video-watching and resting state, using a Graph Signal Processing (GSP) framework. By decomposing the structural graph into Eigenmodes and expressing the EEG activity as an extension of anatomy, GSP provides a way to quantify the structure-function coupling. We elucidate how the structure shapes function during naturalistic tasks such as movie-watching and how this association is modulated by tasks. We quantify the coupling relationship in a region-, time-, frequency-resolved manner. First of all, our findings indicate that the EEG activity in the sensorimotor cortex is strongly coupled with brain structure, while the activity in higher-order systems is less constrained by anatomy, i.e., shows more flexibility. In addition, we found that watching videos was associated with stronger structure-function coupling in the sensorimotor cortex, as compared to resting-state data. Second, time-resolved analysis revealed that the unimodal systems undergo minimal temporal fluctuation in structure-function association, and the transmodal system displays highest temporal fluctuations, with the exception of PCC seeing low fluctuations. Lastly, our frequency-resolved analysis revealed a consistent topography across different EEG rhythms, suggesting a similar relationship with the anatomical structure across frequency bands. Together, this unprecedented characterization of the link between structure and function using continuous EEG during naturalistic behavior underscores the role of anatomy in shaping ongoing cognitive processes. Taken together, by combining the temporal and spectral resolution of EEG and the methodological advantages of GSP, our work sheds new light onto the anatomo-functional organization of the brain.

## 1 Introduction

The functioning of various natural systems is fundamentally constrained by their structure. Analogous to the way the pitch and timbre of music emanating from a flute are intricately tied to the precise design and arrangement of its cylindrical tube, holes, and keys, the functionality of the brain is constrained by its anatomical scaffold (Hagmann et al., 2008; Lynn & Bassett, 2019). However, despite important progress, the precise extent of the association between the brain’s anatomical hardwiring and its dynamic functional properties is not yet fully understood (see Fotiadis et al., 2024 for the collation of current knowledge). Indeed, characterizing the structure-function relationship in neural processes is currently at the forefront of the neuroimaging community’s interest, as illustrated for instance by a recent seminal publication (Pang et al., 2023a) and the various commentary papers it has sparked (Faskowitz et al., 2023; Pang et al., 2023b; Patil et al., 2023).

One line of inquiry that has proven to be promising in investigating structure-function interplay is via spectral graph theory (Spielman, 2007) which provides a framework to express connectivity matrix in graph spectral components using eigenvectors (or eigenmodes) decomposition. In the context of studying the brain, eigen-modes are derived from the Laplacian of connectivity graphs (Bajada et al., 2020), i.e., structural connectome (anatomical links) or functional connectome (statistical dependence between pairs of brain regions). While spectral graph theory analyzes connectivity graphs, an elegant framework that allows to jointly analyze a brain signal onto a connectivity graph is Graph Signal processing (GSP) (Shuman et al., 2013). In GSP, Laplacian eigenvectors are used to decompose the signal in graph Fourier modes by defining a Graph Fourier Transform, thus translating Fourier analysis (and a series of operations such as convolution and filtering) on graphs. GSP is a powerful tool to specifically investigate structure-function interplay and has been successfully applied to analyze brain data in several recent studies (Glomb et al., 2020; Medaglia et al., 2015; Preti & Van De Ville, 2019). The connectivity/GSP framework can be seen as complementary to the classical approach of mapping brain functions in discrete cortical parcellations as it allows for a decomposition of brain activity or structure as a continuum of spectral components (see Lioi et al., 2021 for a perspective review). Crucially, increasing evidence demonstrates the ubiquitous nature of decomposing the connectome, termed as connectome harmonics (Atasoy et al., 2016; Glomb et al., 2021), also called connectivity gradients (Bernhardt et al., 2022) revealing cortical organization (Huntenburg et al., 2018; Margulies et al., 2016), how anatomy shapes the functional Magnetic Resonance Imaging (fMRI) (Preti & Van De Ville, 2019), the Electroencephalography (EEG) (Glomb et al., 2020), and the Magneto-electroencephalography (MEG) (Griffa & Preti, 2022).

Pioneering studies with resting-state fMRI data have investigated macroscale brain organization and found that the first connectivity gradient reveals a hierarchy spanning between unimodal (sensory systems) to transmodal (higher-order systems) association regions (Margulies et al., 2016). To study how the structure-function coupling is organized in the brain, Preti and Van de Ville (Preti & Van De Ville, 2019) developed a metric from GSP called the Structural-Decoupling Index (SDI). The term *Coupling* in this context refers to brain activity that closely matches the underlying structure, whereas the *decoupling* refers to brain activity less tied to the structure. In particular, the structural connectome imposes constraints uniformly across all its harmonics, influencing both the excitation of Eigenmodes with long spatial wavelengths (lower Eigenmodes) and those with short spatial wavelengths (higher Eigenmodes). The high Eigenmodes and the low Eigenmodes characterize the localized activities, and the broad-scale activities respectively. Activities expressed by lower harmonics, that propagate through the cortex via anatomical structures, are seen as ‘coupled’ to these structures. Conversely, localized activities, which signify flexibility and lesser dependence on the underlying network, are described as ‘decoupled’ from these structures (see Medaglia et al., 2018 for an elaborative interpretation). Thus, SDI essentially quantifies the ratio of structurally decoupled activity over the coupled activity. Preti and Van de Ville investigated the dependency of fMRI signal on the underlying structural connectome estimated with Diffusion-Weighted Imaging (DWI) during rest and found a similar hierarchy revealed by Margulies et al., 2016, with the coupling of fMRI activity gradually decreasing along the unimodal-transmodal axis (Preti & Van De Ville, 2019). While the majority of the contributions in this line of work has used MRI imaging, a handful of studies have attempted to characterize the dependency of the electrophysiological signals on the structural connectome. For instance, SDI has been used to quantify the relationship of fast temporal activity with MEG in Griffa and Preti, 2022, indicating a different gradient of organization as compared with fMRI, with coupled sensory areas (task-positive) and decoupled default mode network areas (task negative). Other recent studies investigating EEG visual and auditory event-related potentials found that during these tasks, the cortical activity can be compactly represented by a sparse set of structural connectome harmonics (Glomb et al., 2020; Rué-Queralt et al., 2021, 2023). Structure-Function coupling of the EEG has also been studied during epileptic seizures (Rigoni et al., 2023) with results showing that epileptic activity is more coupled to a generic structural graph during the spike. These studies focused on characterising the interplay between anatomical structure and EEG activity in well isolated and/or evoked events. This is in contrast with the works applying the GSP framework to fMRI which have primarily focused on resting state. Moreover, to the best of our knowledge, all the brain studies using the gradients/GSP framework have considered a common structural graph, estimated from either unrelated healthy subjects (such as in Griffa and Preti, 2022; Preti and Van De Ville, 2019) or as the average connectivity matrix across the population (Rué-Queralt et al., 2023). Taken together, despite the important progress in probing the nature of structure-function coupling through the lens of task-based electrophysiological responses, the relationship between the continuous EEG and the underlying structural connectivity remains elusive.

This study tackles for the first time the fundamental question of the relationship between continuous EEG activity and individual structural connectomes using GSP. We estimate EEG cortical activity during rest and video watching and use the Laplacian eigenmodes of the individual structural graph to decompose the source-localized activity into graph-informed components. We then estimate the SDI metric to characterize the structure-function coupling relationship of EEG in different frequency bands. Precisely, we address the following open questions: a) How is the continuous EEG during rest and video watching constrained by the underlying individual structural connectivity? b) Are the rest and video-watching EEG activities constrained by the structure in the same way? c) Does the structure-function coupling exhibit different patterns for different EEG frequency bands? d) Does the function depend on the structure similarly throughout the entire period of Video ? Using high quality EEG and individual DWI data from a large open access dataset and rigorous validation/estimation of GSP derived metrics, we provide novel insights into how the anatomy shapes the continuous electrophysiological activity of the human brain, and how this relationship is modulated by behavior.

## 2 Methods

### 2.1 Data and Preprocessing

We analyzed a subset of the EEG and DWI data acquired by the Healthy Brain Network (HBN) (Langer et al., 2017). In particular, we used high-density EEG recorded using a 128-channel Geodesic Hydrocel system at a sampling frequency of 500 Hz while subjects were (i) resting with open eyes (100 sec), (ii) watching a clip from Despicable Me 2 (Video 1 ^1^, 170 sec) and (iii) watching a clip from Fun with Fractals (Video 2 ^2^, 162 sec). For the purpose of test-retest reliability analysis, we considered Video 2, which is a tutorial-oriented audiovisual stream containing fractals, as compared to the clip from the animation movie with people inside (Video 1). The released EEG data includes preprocessing steps such as electrode data quality checks, notch-filter at 60Hz, high-pass filter at 0.1Hz and artifact signal correction (more details in Langer et al., 2017). In addition, we excluded 27 electrodes and only selected subjects having a good EEG quality determined as part of the quality assessment in Nentwich et al., 2020. We identified a group of 43 subjects (aged 14 - 21.8) based on the following criteria: i) age of at least 14 years; ii) having both resting state and video watching EEG; iii) undergone DWI scans in order to estimate the subject-specific structural connectome.

### 2.2 EEG Source Reconstruction

We used Freesurfer (Fischl, 2012), Boundary Element Method (BEM), and exact Low-Resolution Electromagnetic Tomography (eLORETA) (Pascual-Marqui et al., 2011) to estimate the cortical sources. To define the cortical surface and cortical mesh, we used Freesurfer’s fsaverage version 5. We characterized the electromagnetic properties of different brain segments using BEM and used the consensus montage of the 128-EEG channel array while computing the forward operator. We used subject-specific eyes-closed and eyes-open resting state (20 sec) to compute the noise-covariance matrix for estimating the cortical sources for resting-state eyes-open and video-watching EEG respectively. We computed the inverse solution using eLORETA (Pascual-Marqui et al., 2011), a minimum-norm estimate algorithm as implemented in MNE-Python (Gramfort et al., 2013). We estimated EEG cortical sources for the entire time series on the 20484 vertices of the cortical mesh, and then parcellated the full-length signal into 360 Regions of Interest (ROIs) of the HCP-MMP atlas (Glasser et al., 2016). We bandpassed the full-bandwidth EEG signal into the standard frequency bands: *θ* (4 - 8Hz), *α* (8 - 13Hz), Low *β* (13 - 20Hz), High *β* (20 - 30Hz), and *γ* (30 - 40Hz). To this end, we first downsampled the raw signal to 125Hz (anti-aliasing filter; performed using mne.epochs.resample()), followed by applying a fifth-order butterworth filter (implemented in scipy.signal.butter()). We then applied the Hilbert Transform to extract the signal envelopes for the narrow-band signals (implemented in scipy.signal.hilbert()). We then assessed the structure-function relationship for two types of EEG signals: 1) EEG raw signal; 2) the narrow-band cortical EEG envelopes in the different frequency bands across different conditions (Video 1, Rest, and Video 2). We also quantified the structure-function association for the bandlimited signals, and the Short-time Fourier Transform (STFT) coefficients of the cortical EEG (supplementary material, section 11).

### 2.3 Structural graph

#### 2.3.1 Connectome reconstruction

We used Qsiprep (Cieslak et al., 2021) to process the structural (T1w) and DWI images in the Brain Imaging Data Structure (BIDS) format. The preprocessing pipeline included head motion correction, susceptibility distortion correction, MP-PCA denoising, coregistration to T1w images, spatial normalization using ANTs, and tissue segmentation (Cieslak et al., 2021). We used the MRtrix3 (Tournier et al., 2019) pipeline available through Qsiprep for fiber reconstruction. In particular, we employed multi-shell multi-tissue constrained spherical deconvolution (mrtrix multishell msmt ACT-hsvs) for estimating the fiber orientation distribution using the Dhollander algorithm (Dhollander et al., 2019). We then applied tckgen, specifically iFOD2, a probabilistic tracking method that generates 10^7^ streamlines, with the weights for each streamline calculated using SIFT2 (Smith et al., 2015). We set the T1w segmentation reconstructed through Freesurfer as an anatomical constraint. We used the same HCP-MMP atlas for fiber segmentation and we quantify the structural relationship between pairs of ROIs as the density of the fibers calculated as the sum of the fibers connecting two regions divided by the region volumes as in Preti and Van De Ville, 2019. The estimated individual connectomes were then used to assess subject-specific structure-function coupling.

#### 2.3.2 Eigendecomposition of the connectome

Structural connections between different brain regions can be effectively modeled as graphs (Bassett & Sporns, 2017; Lynn & Bassett, 2019) (Fig 1a), and the spectral properties of the graphs can be further studied using the Laplacian operator (Spielman, 2007). Let us consider an undirected weighted graph *G* =*< V*, ℰ, *W >*, where *V* is a set of *N* elements called vertices and ℰ ⊂ *V* × *V* the set of edges connecting unordered pairs of vertices with scalar weights *w*. In this work, the nodes correspond to HCP-MMP ROIs, and the edges represent white matter fiber links between those nodes weighted by the strength of the connections (e.g. fiber density). The degree matrix **D** of a graph *G*, is a diagonal matrix such that 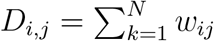 and 0 otherwise. The Adjacency Matrix **A** is a square matrix of dimension *N* × *N* in which each element is different from zero only if the corresponding edge exists. Given **A** and **D** we can define a Laplacian operator. In line with previous studies such as Preti and Van De Ville, 2019, we used the symmetric normalized Laplacian, which is defined as **L** = **I** - **D**^−**1***/***2**^ **AD**^−**1***/***2**^, where **I** is identity matrix. **L** the Laplacian is a real symmetric matrix and thus can be diagonalized as

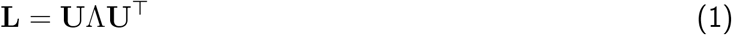

where **U** denotes the base of orthogonal eigenvectors and **Λ** the diagonal matrix of eigenvalues, sorted in increasing order. Eigenvectors (often referred to as eigenmodes) capture the spatial frequencies of the graph and are ordered by an increasing level of spatial frequency (i.e., low frequency to high frequency). In particular, low-frequency eigenmodes capture the main axes of information diffusion in the network while high-frequency eigenmodes are more scattered, as illustrated in Fig 1b.

**Figure 1:**
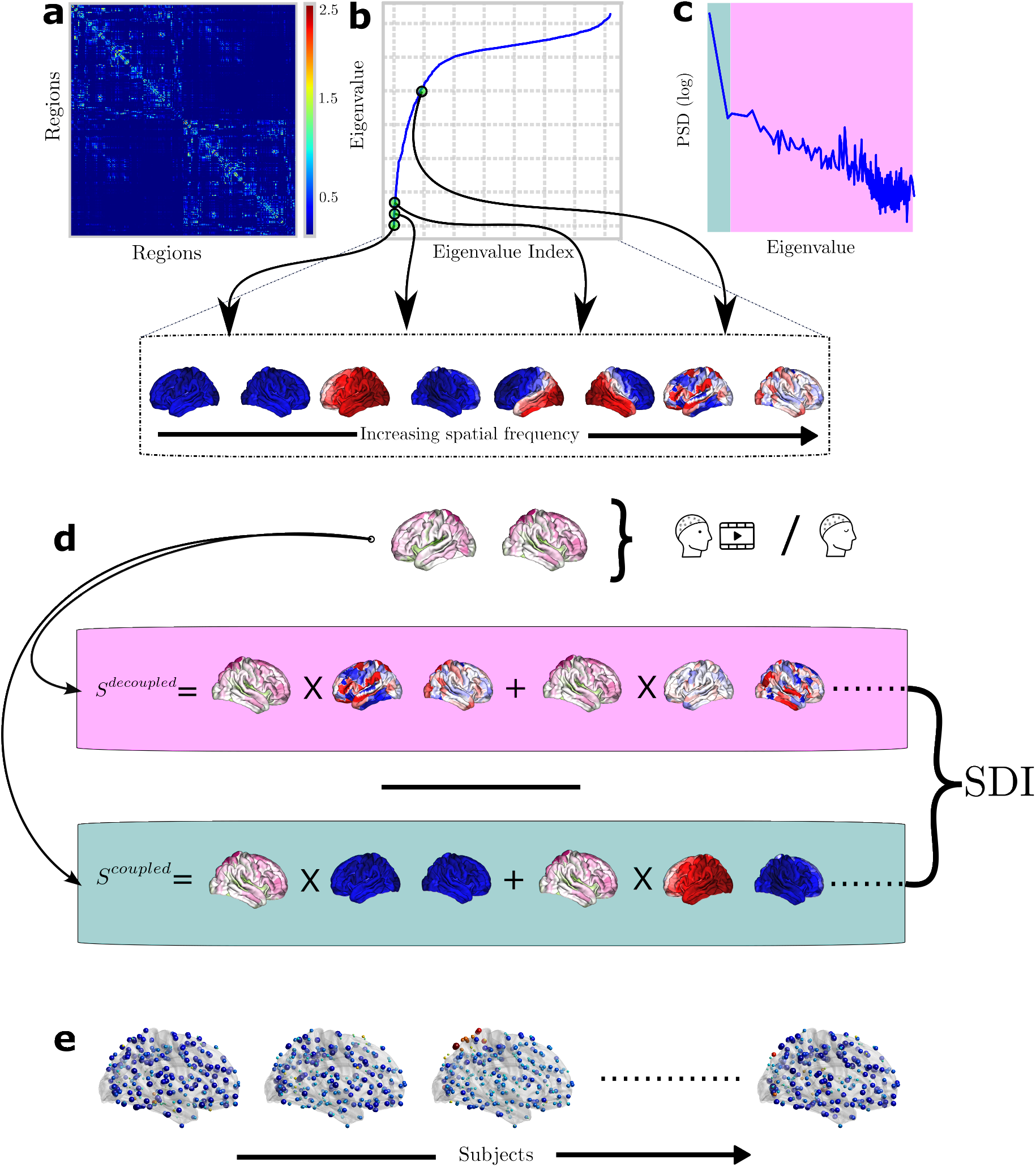
Illustration of the methods pipeline. **a** The structural connectome is estimated for each individual (here we show the consensus structural graph in natural logarithmic scale). **b** The spectral decomposition of the structural connectome Laplacian results in eigenmodes that capture the spatial harmonics progressing from the smoothest (first eigenmode, corresponding to the lower part of the spectrum) to high frequency eigenmodes, corresponding to higher part of the spectrum. Eigenvectors are orthogonal to each other, so the EEG activity can be expressed as a weighted linear combination of them using the Graph Fourier Transform. **c** Averaged Power Spectral Density (PSD) of EEG graph signals (*ℓ*2-normed) revealing the power distribution of the source EEG on the graph. Median power split criterion was used to split the graph power spectrum in order to identify the low/high-band pass filters cut-off frequency. **d** EEG cortical signals are high and low pass-filtered using graph filters to isolate the activity which is respectively decoupled and coupled to the underlying structural graph. The SDI index is calculated as the ratio between the decoupled and coupled activity (Preti & Van De Ville, 2019). **e** SDIs for a few representative subjects.

#### 2.3.3 Graph Fourier Transform and Structural Decoupling Index estimation

The set of scalar values residing on the vertices of the graph is called graph signal. In this work the graph signal *S*_*t*_ is the cortical EEG at time t associated with each node (ROI) in the graph. Eigendecomposition provides a Fourier basis for the graph signal, enabling the application of classic signal processing techniques such as Fourier transform or filtering. The Graph Fourier Transform (GFT) transforms the signals defined on the vertices of a graph into the graph spectral domain

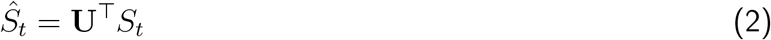

The original signal *S*_*t*_ at time t can be reconstructed back using Inverse GFT (equation 3).

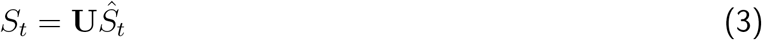

Once we have the notion of a graph Fourier transform, we can generalize from classic Fourier domain to define frequency filtering, or graph spectral filtering such as 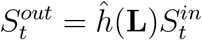 where

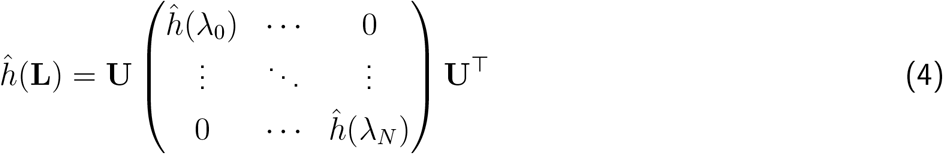

We can therefore isolate the low frequency and high frequency components of the graph signal with a graph low-pass and high-pass ideal filters,

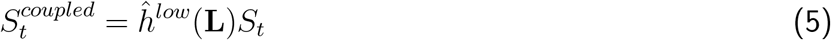

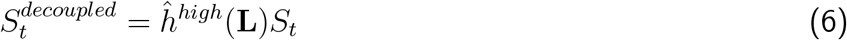

where 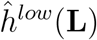 and 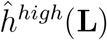 act as band-pass filters, extracting the low- and high-frequency spatiotemporal activity denoted as 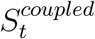 and 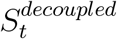. Specifically, 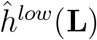 retains only the first *C* diagonal elements and pads the rest with zeros while 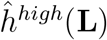 keeps only the diagonal elements corresponding to the last *N* – *C* eigenvalues. Notably, 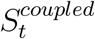 is the EEG activity that is closely aligned to the structural graph, while the high graph frequency activity 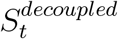 is less constrained by the underlying network (Lioi et al., 2021). The Structural-decoupling index (SDI) (Preti & Van De Ville, 2019) is defined as the ratio between the norm (*ℓ*1-norm) of 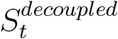 and 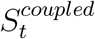. We applied binary logarithm as in Preti and Van De Ville, 2019, thus the positive and negative SDI values indicates decoupling and coupling, respectively. We computed SDIs for each region across all subjects and the resulting spatial maps are presented in Fig 1e for a few subjects. As in Preti and Van De Ville, 2019, we identified as cut-off frequency between low and high frequencies the one that splits the spectrum into two parts with equal energy (median-frequency)(Fig 1b and c). The cut-off frequency varies among subjects but its median is the same across conditions (Video 1, Rest and Video 2: median *C* = 2 for the raw EEG signal). Readers can consult the supplementary material (section 10) for a different strategy to compute the cut-off frequency.

We investigated the time-varying aspect of SDI. To this end, we adapted the SDI metric. In particular, we considered the entire time series into ”epochs” of length 1s and computed the cut-off frequency C in a temporally localized manner. This results in *C* cutoff adapted to each time window. Furthermore, the *ℓ*1-norm is applied over the 1s window of 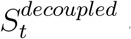 and 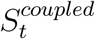 component. Thus, applying the ratio on the norms results in SDI quantified at the resolution of 1Hz.

### 2.4 Decoding of the SDI maps

To establish the functional relevance of SDI maps, we performed a NeuroSynth (Yarkoni et al., 2011) meta-analysis similar to previous studies (Margulies et al., 2016; Preti & Van De Ville, 2019). First, unthresholded SDI maps were segmented into bins with a 10%ile increment and binarized. Next, we used ROIAssociationDecoder function in NiMare to decode the 10 binary masks using the Neurosynth database. This database contains over 500,000 activation maps from over 14K studies (more details in NiMare website and in Salo et al., 2022). ROIAssociationDecoder synthesizes this entire database and perform spatial correlation between input masks and the meta-analytic maps to extract the relevant topic-terms. We then transformed the resulting correlation coefficient into z-statistics and showed significant results corresponding to a p-value*<*0.001. Note that unlike the previous studies that performed decoding (Margulies et al., 2016), for an easier comparison between task conditions and with the previous studies, we manually ordered the topics based on the hierarchy in lower order to higher order systems.

### 2.5 Null models and statistical analysis

Firstly, we assessed the level of statistical significance of SDI maps. To do this, we compared the empirical SDI with graph-informed surrogate SDI, following the methodology to obtain null models outlined in Preti and Van De Ville, 2019. In particular, to generate graph-informed surrogates, we employed spectral randomization of the graph (Pirondini et al., 2016) that randomizes the phase of the GFT coefficients, while preserving the auto-correlation of the graph Laplacian.

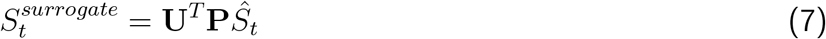

Here, **P** represents a diagonal matrix with random +1 and -1 enforcing randomization for the inverse GFT to produce the surrogate cortical signal. A total of 19 surrogates were generated, maintaining *α* = 1*/*(1 + 19) = 0.05. We computed surrogate SDI for each condition (Video 1, Rest, and Video 2) using the surrogate EEG time series, in the same way as for the empirical SDI. The surrogate SDIs served as a noise floor against which individual empirical SDIs were tested. We then assessed consistency across subjects using a threshold determined based on binomial cumulative distribution with the following parameters: 43 subjects, 100 trials, the rate of success at p=0.05 (Preti & Van De Ville, 2019). We extended this pipeline to perform group-level SDI over time.

We also assessed differences between video-watching and rest with a paired statistical test between the empirical SDIs. Within each EEG frequency band, we performed a t-test between the conditions followed by FDR-correction (*α* = 0.05).

Finally, we assessed the difference in structure-function coupling patterns across EEG frequency bands. To do this, we aggregated the SDIs from HCP-MMP ROIs into Yeo-Krienen networks (Yeo et al., 2011) and computed Spearman’s *ρ* between pairs of frequency bands.

### 2.6 Test-retest reliability with Video 2

To test the robustness of the findings presented for Video 1, we performed the same analysis for Video 2. We used intra-class correlation coefficient (ICC) as the reliability index (Koo & Li, 2016). The ICC estimates and the 95% confidence intervals were computed using the Pingouin package in Python. Specifically, we used the two-way random-effects model of ICC to determine the inter-rater reliability between two videos, treating the Video 1 and 2 as independent raters. The choice of the random-effects model for ICC accommodates potential sources of variability that could influence EEG recordings, including different days, variations in equipment, and different experimenters.

## 3. Results

In what follows, we assess how anatomy shapes the source-localized EEG across naturalistic tasks, besides Rest. We quantify the relationship in a region-, time-, and frequency-resolved manner.

### 3.1 Brain-wide coupling and focal decoupling patterns across video and rest

We began by investigating the structure-function coupling of the raw EEG signals during video-watching and the resting-state. First, we show in Fig 2a global brain activity patterns during Video and Rest, using averaged spatial map of the source-localized raw EEG across subjects and time. Resulting spatial maps were then zscored to highlight the salient regions. These two maps summarize the brain activity over the entire period of video and rest, and show average brain-wide activation that is mostly similar between conditions. As our focus is investigating structure-function coupling, and due to the continuous nature of the signals, we did not seek to compare brain activity statistically.

**Figure 2:**
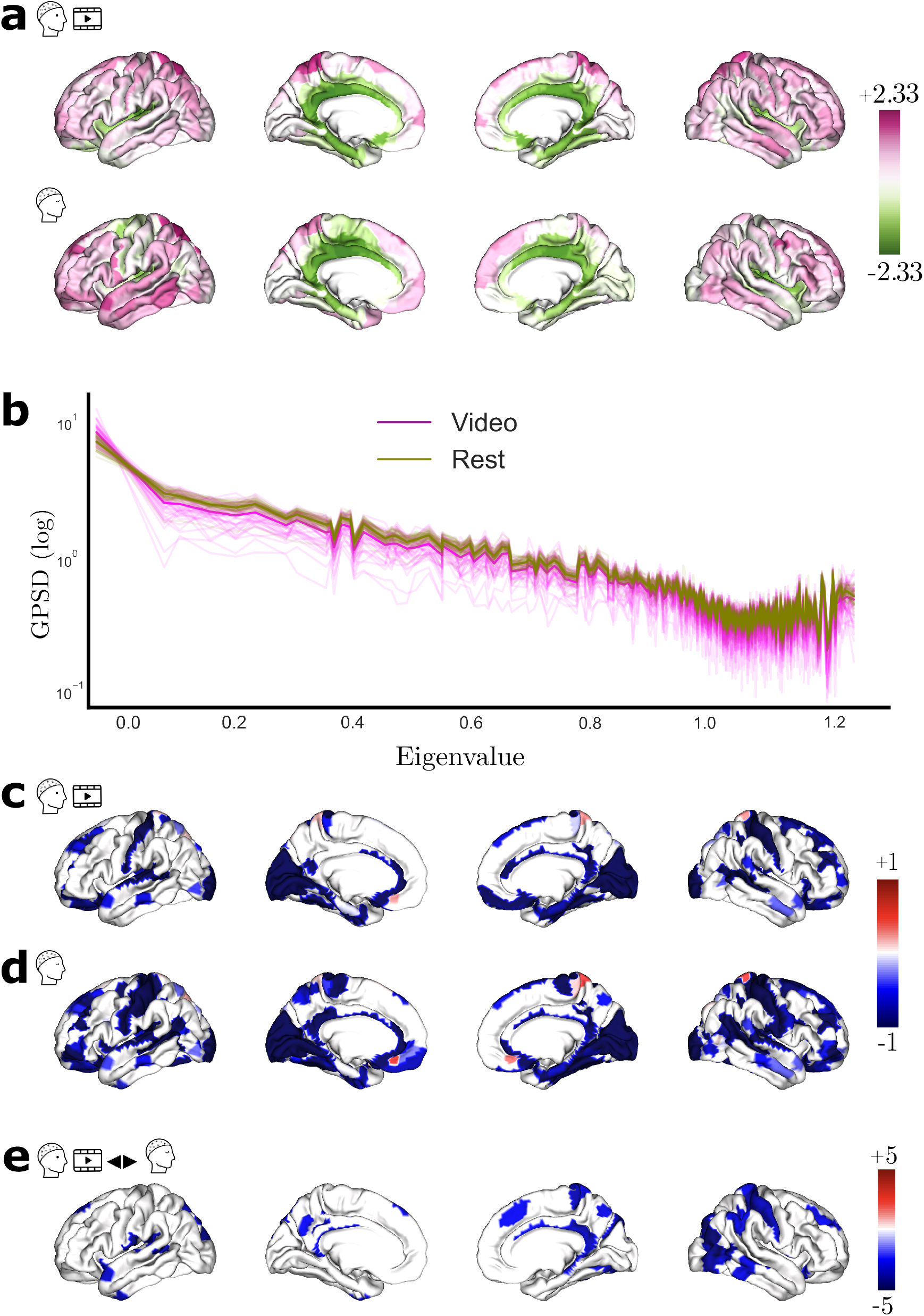
Source-localized EEG and SDI maps. **a** Averaged cortical activity (in zscore) for Video 1 (top row) and resting-state. **b** Averaged graph PSD for all subjects (magenta - Video 1; olive - Rest). The power spectrum is in log scale, displayed as a function of consensus structural graph eigenvalues. **c** Group level SDI maps (using subject-specific structural connectome) for Video 1 and **d** for Rest. For visualization purposes only, and due to the log scale of SDI, values are clipped to -1 / +1. **e** SDI contrast t-values maps (FDR corrected) resulting from the statistical comparison between Video 1 and Rest, with blue (and red) denoting stronger (and weaker) coupling during the movie than during rest.

Average graph PSD for the video-watching and resting state cortical signal are presented in Fig 2b for video (in magenta) and rest (in olive). To calculate the average graph PSD across subjects we considered the consensus graph. The graph PSD of the cortical activity has a 1/f-like shape, with the largest part of the power in the low frequencies. This indicates that the low-frequency connectome harmonics carry dominant role in explaining the given cortical signal.

For the rest of the results, we quantified the structure-function relationship using individual connectomes. The group-level SDI map during video-watching is represented in Fig 2c. This map, thresholded using group-level significance (see section 2.5) highlights the coupling (blue) and decoupling (red) of the electrophysiology on the anatomical scaffold. Video-watching activity is mostly coupled to the structure across the whole brain. Regions exhibiting the strongest coupling are the Primary Somatosensory cortex (Area 2; SDI: -1.8), Primary Sensory cortex (SDI: -1.8) and Second Visual area (V2; SDI: -1.8). Activities from the regions such as the Parahippocampal Area 2 (SDI: 0.62), Inferior Frontal Sulcal area (IFJp; SDI: 0.5), and Posterior parietal cortex (Area 5L; SDI: 0.2) are decoupled from the underlying anatomy.

The group-level SDI map for the rest condition is displayed in Fig 2d. During resting-state, EEG is mostly coupled to the structural graph, similarly to the video. Notably, primary somatosensory cortex (BA1, SDI: -1.78), Visual V2, and V3 (SDI: -1.74 each) exhibited strong coupling, whereas the Parahippocampal area 2 (SDI: 0.71), Inferior Frontal Sulcal area (IFJp; SDI: 0.6), and Subgenual Anterior Cingulate cortex (Area s32; SDI: 0.38) showed decoupling from the structure. While some effects appear to be specific to video-watching tasks, most patterns appear to be consistent between rest and video-watching.

### 3.2 Stronger coupling to the structure while watching movie as compared to rest

To further investigate the differences between video and resting state, we compared the SDI in the two conditions (Fig 2e). The statistical comparison (Paired t-test, FDR-corrected at *α* = 0.05) revealed differences in the coupling relationship between video and rest. Note that these are to be interpreted as weaker coupling (red), and stronger coupling (blue) during movie-watching relative to resting state. Regions that exhibit stronger coupling during video watching as compared to rest were found to be widespread across the whole brain. In particular, stronger coupling was observed in the Visual area 7 (V7; t(42) = -5), Inferior parietal cortex (Area PGP; t(42) = -4.65) and Middle temporal area (t(42) = -4.64) during movie-watching. No region exhibited weaker coupling. Raw EEG is tethered consistently within condition during video and the relationship is strengthened during movie. Given this observation, one may wonder whether the coupling relationship of the EEG frequency bands could exhibit a distinct behavior.

### 3.3 Sensory regions show temporal stability in Structure-Function association

We harnessed here one of the advantages EEG offers to study the structure-function association in a temporally fine-grained manner. We computed the SDI over time for raw EEG signal, the group-level time-resolved SDI map is presented in Figure 3a. We can observe that while the activity in most regions is coupled to the structure, activity in a few regions is decoupled, and this trend is rather consistent over time. Interestingly, there does not seem to be any spatial variability in the SDI, i.e. we cannot clearly see a ROI that switches from coupling to decoupling or vice versa. This coupling (or decoupling) appears to remain largely stable over time. To investigate which regions exhibit particular stability or relative fluctuation over time, we considered analyzing the SDI in a region-resolved manner. We considered standard deviation as a measure to index the temporal fluctuation of the SDI, and the associated results are presented in Figure 3bc. We show the regions that exhibit lowest (highest) temporal fluctuation, characterized by bottom (top) 20%ile. The corresponding results are presented in the top and bottom row respectively. We observed that the coupled regions such as sensorimotor and Posterior Cingulate Cortex (PCC) exhibit relatively low fluctuation over time (pink), and the regions that are decoupled to the structure are associated with higher temporal fluctuation. We replicated the findings with Video 2 (Figure 3c)

**Figure 3:**
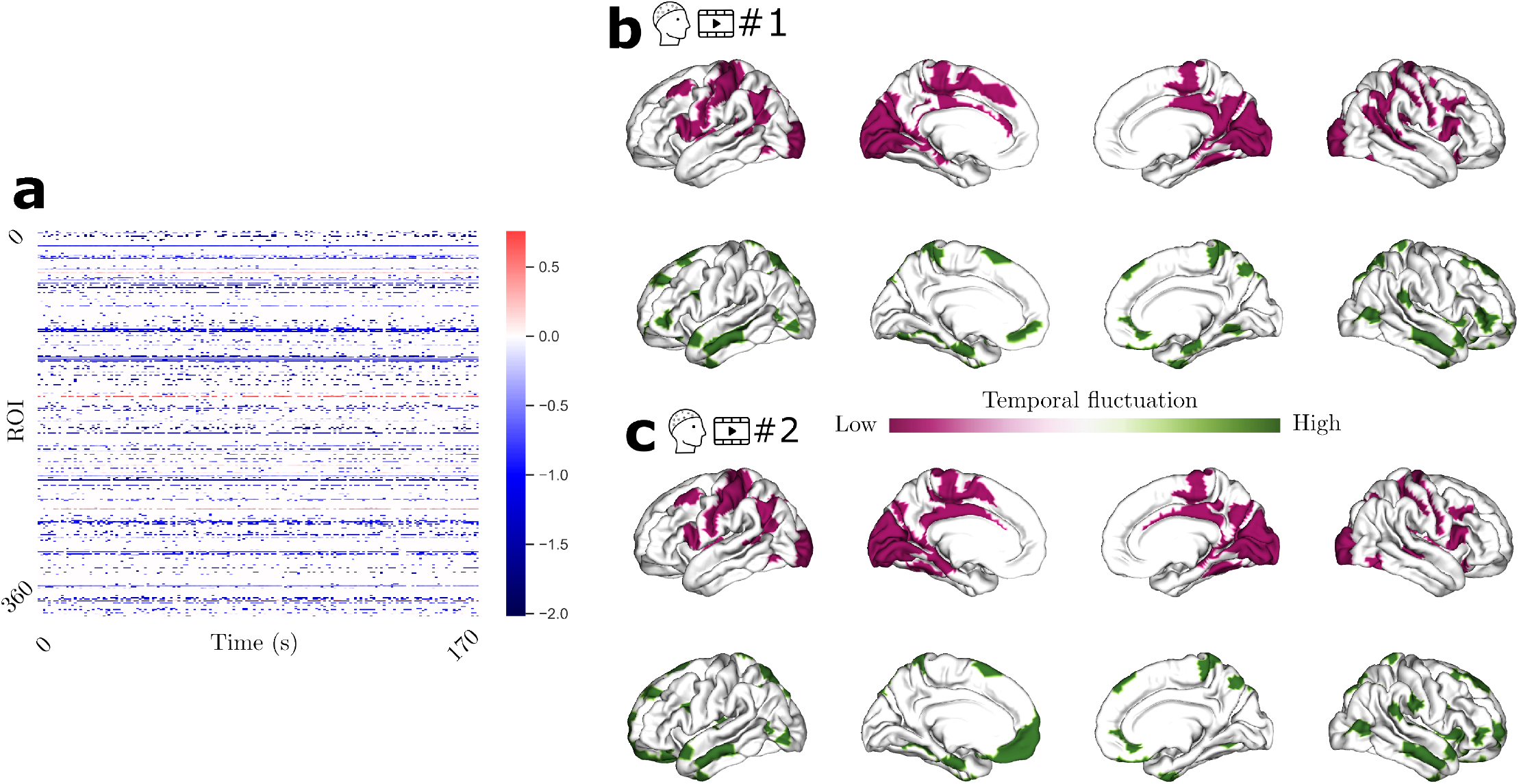
Time-resolved SDI for the raw EEG. Panel a) depicts the group-level SDI over time. The top and bottom row depicts the regions that exhibit lowest (bottom 20%ile; pink) and highest (top 20%ile; green) temporal fluctuation respectively for Video 1 and 2 (panel b and c).

### 3.4 Structure-Function relationships are similar across EEG frequency bands

We investigated whether all EEG frequency bands (cortical envelopes) are related to the underlying anatomical structure in the same way through an analysis focused on each of the EEG frequency bands i.e *θ* (4 - 8Hz), *α* (8 - 13Hz), Low *β* (13 - 20Hz), High *β* (20 - 30Hz), *γ* (30 - 40Hz). Results are presented in the first four columns of Fig 4. Upon closer examination of the SDI values specifically for the regions such as the Retroinsular cortex (Mean SDI: -1.2, *σ*: 0.1), Primary Auditory Cortex (Mean SDI: -1.1, *σ*: 0.2) exhibited consistent coupling, whereas the Parahippocampal area (Mean SDI: 0.8, *σ*: 0.07) exhibited consistent decoupling across frequency bands. We compared the coupling relationship between video and rest for the specific EEG frequency bands.

**Figure 4:**
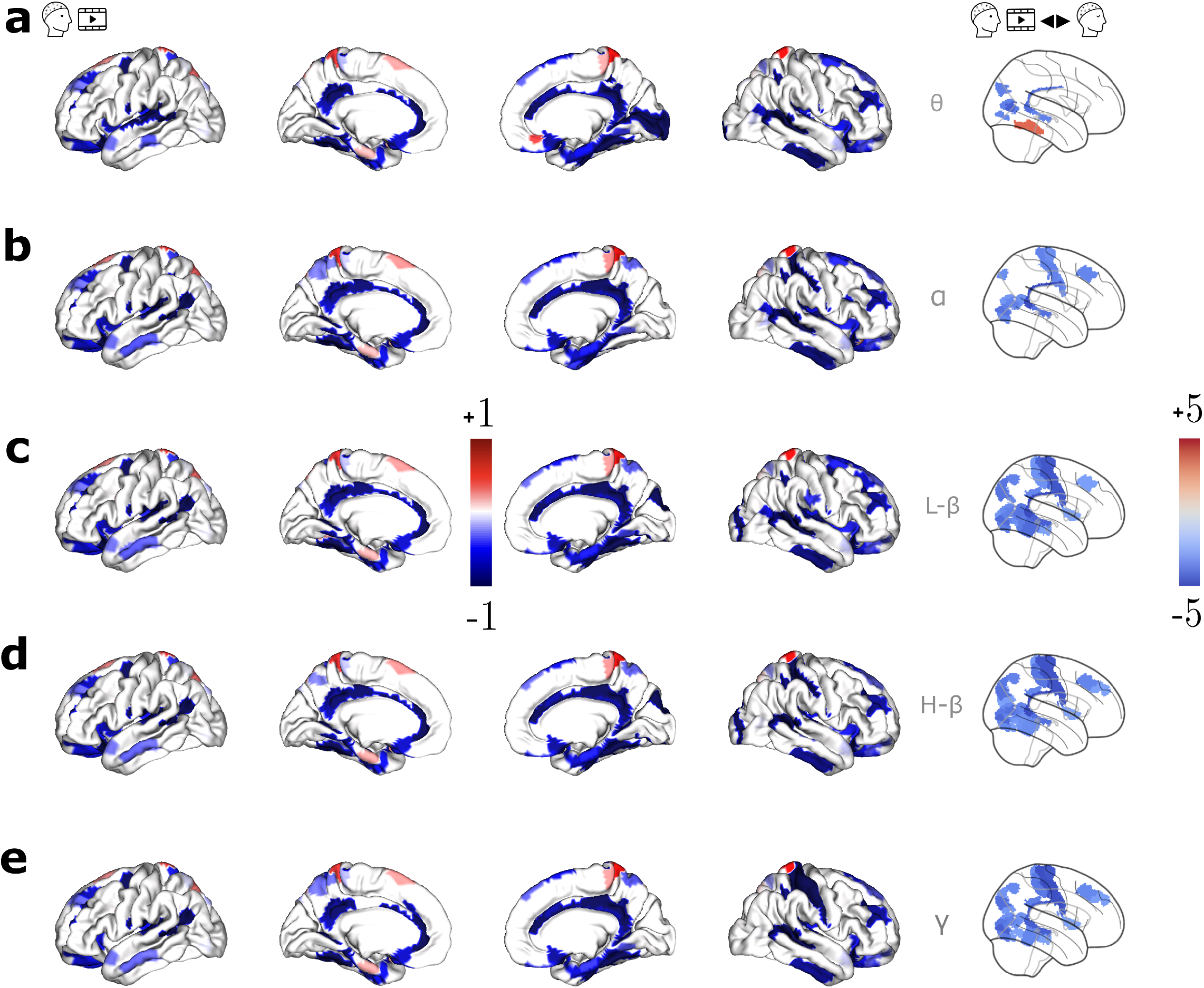
Group SDI maps during video watching (Video 1) in different EEG frequency bands: **a** *θ* (4-7Hz), **b** *α* (8-13Hz), **c** Low *β* (13-20Hz), **d** High *β* (20-30Hz), and **e** *γ* (30-40Hz). The first four columns display group-level significant SDI maps, with red and blue indicating respectively decoupling and coupling. The last column shows the group-level SDI contrast t-values maps (FDR corrected) resulting from the statistical comparison between Video 1 and Rest, with red (and blue) denoting weaker (and stronger) coupling during the movie than during rest

Results (presented in the last column of Fig 4) revealed that the stronger coupling observed during movie-watching in areas such as Retrosplenial complex and Motor cortices still holds for different EEG frequency bands. We quantified the structure-function association for the bandlimited signals and the STFT coefficients (supplementary material, section 11), and found a similar topography across frequency bands.

We further investigated how the SDIs are related across frequency bands by computing the Spearman’s correlation coefficient in 7 Yeo-Krienen networks. Results showed a significant correlation across all EEG frequencies and in all seven networks (average *ρ* = 0.93, all p-values *<* 10^−7^).

### 3.5 SDI spatial maps are consistent across two different videos

To ensure the reliability of the findings we investigated the structure-function association by swapping the video available in the HBN dataset. To this end, we performed a test-retest reliability of our study’s findings with Video 2. We performed a reliability analysis on a mean rating (raters corresponding to the two videos) with a two-way random-effects model. Our analysis revealed that the ICC estimates is 0.83 and the 95% CI is [0.79, 0.86] for the said parameters, indicating a good degree of reliability across videos. Furthermore, we ran a reliability analysis on the SDI contrasts between rest and Video (rater 1 = Video 1 vs Rest; rater 2 = Video 2 vs Rest). Our findings revealed that the ICC estimates were 0.68, and the 95% CI was [0.6, 0.75], indicating a good amount of reliability in contrast between video and rest. We also assessed the extent of agreement between task conditions by computing ICC coefficients for either of the videos and rest. The results revealed that the ICC was 0.79 and the 95% CI was [0.75, 0.83] when comparing video 1 with rest and the ICC was 0.78 and the 95% CI was [0.73, 0.82] with the video 2. Taken all the pairs of ICC tests together, we can observe the agreement between videos was higher than the agreement between distinct task conditions, suggesting the topography is more similar between videos than between videos and rest.

Fig 5 shows SDI maps corresponding to video 2 (Fig 5a), and contrast between video 2 and resting state (Fig 5b), both FDR-corrected at *α* = 0.05. As with Fig 2e, the blue tail in Fig 5b indicates stronger coupling during video 2 compared to the resting state. Our results revealed that the regions such as Visual areas (V3A, V6A; t(42) = -5.4, -5.4 respectively) Superior parietal lobule (Area 5m; t(42) = -5.2) exhibited stronger coupling during movie. Overall, the trend was largely similar to the results obtained for Video 1 (Fig 2e).

**Figure 5:**
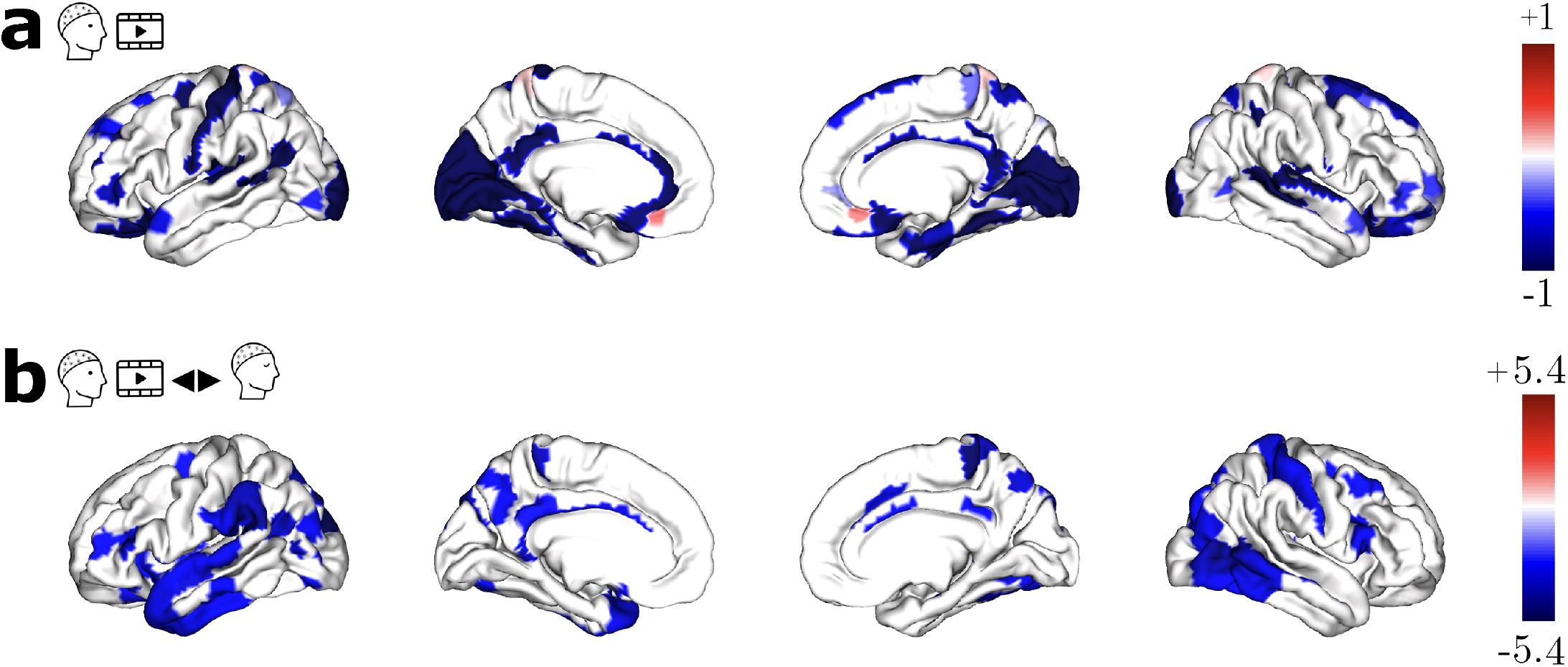
Structure-Function relationship for Video 2. **a** Group-level SDI map for Video 2. **b** SDI contrast t-values maps (FDR corrected) resulting from the statistical comparison between Video 2 and Rest, with red (and blue) denoting weaker (and stronger) coupling during the movie than during rest

### 3.6 Functional decoding of the structure-function relationship

We performed functional decoding of SDI spatial maps with a meta-analytic search of brain maps and associated topics terms in the literature. The results for decoding the raw SDI maps are presented in Fig 6. Panels a) and c) show the results for Video 1 and 2, while panel b) shows the results for the resting state.

**Figure 6:**
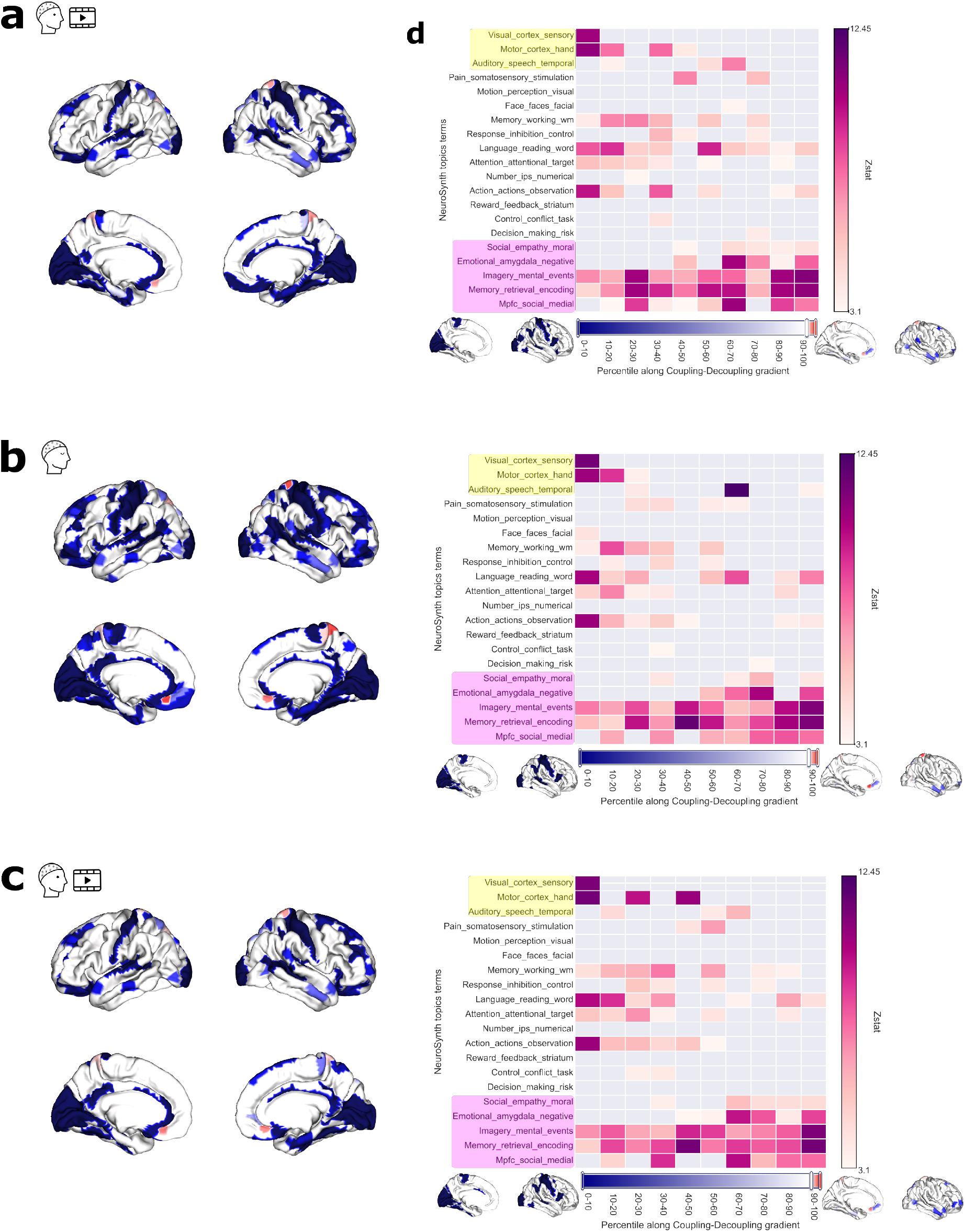
Results of the Neurosynth meta-analysis of SDI maps for **a** Video 1, **b** Rest and **c** Video 2. Decoding results, shown as thresholded z statistics as a function of topic terms, are reported on the right as a function of percentile along the decoupling index gradient (i.e. binned SDI values). Yellow and pink topic terms correspond to unimodal and transmodal systems respectively, following the convention used in Preti and Van De Ville, 2019. The unthresholded maps, corresponding to first and last decile bins, used to perform decoding are presented on the sides of the colorbar. The color of the horizontal colorbar corresponds to coupling in blue, and decoupling in red. The corresponding thresholded group-level SDI spatial maps are reported on the left.

The thresholded spatial maps are shown in the first column as a reference and the second column displays the decoding results. The unthresholded spatial maps that were used to perform decoding are presented in both sides of the colorbar. The arrangement can be thought of as a strong coupling-weak coupling gradient, where the ascending order indicates more untethering from the underlying structure. SDI maps were binned at 10%ile increments, and decoding was performed for these 10 ”activations maps”. The lower percentiles indicate strong coupling to the underlying structure (corresponding spatial map on the left side of the colorbar), whereas the higher percentiles correspond to weaker coupling (corresponding spatial map on the right side of the colorbar). Our decoding analysis of the spatial map for both Video 1 and Rest revealed that the unimodal visual systems and motor systems were associated with strong coupling (0 to 40th percentiles in Fig 6a and 6b, topics highlighted in yellow). Though the other unimodal system Auditory is associated to coupling, this system can be observed to be relatively flexible extending between 20th to 90th percentile. Results obtained for Video 2 (Fig 6c) indicate a closer similarity to Video 1 than to rest, with the key difference being that during rest the correlation of the auditory system was also found around the coupling-decoupling divergence. However, higher-order systems such as mental imagery, emotions, and memory retrieval / encoding (Fig 6, topics highlighted in pink) were more distributed across the strong coupling-weak coupling gradient, similarly in Video 1, Rest and Video 2.

## 4 Discussion

The static anatomical scaffold in the brain gives rise to complex and dynamic functions ranging from primary perceptual tasks to higher-order cognitive processes (Honey et al., 2009; Park & Friston, 2013; Pascucci, Rubega, et al., 2022; Tewarie et al., 2019). Here, we studied how brain activity is coupled to the underlying anatomy, by analyzing how the source-localized EEG is linked to structural connectivity (derived from individual DWI data) during movie-watching and during rest. A Graph Signal Processing framework (Preti & Van De Ville, 2019) was sought out to quantify structure-function coupling. In particular, the eigenvectors of individual structural graph Laplacians were used to define a Graph Fourier Transform and isolate low and high graph-frequency components. The association between structure-function was then quantified using the SDI, calculated as a ratio between low- and high graph-frequency components of the brain activity (Preti & Van De Ville, 2019).

The Laplacian matrix is central to our analysis. Laplacian is related to the heat equation, with the laplacian decomposition reflecting a diffusive process which has locality-preserving properties capturing the intrinsic geometry (Belkin & Niyogi, 2003). Diffusion-based models are a kind of several other canonical propagation models such as random walks, which reflect a stochastic diffusion across the network (Zamora-López & Gilson, 2024). There are several implications in terms of graph spectral properties, and how it characterizes the brain dynamics. In the Laplacian matrix, the diagonal elements represent node degrees, which can be interpreted as ”heat sources.” The heat (or neural activity) then diffuses across the network to connected nodes, with the diffusion weighted by the strength of the structural connections in the brain’s connectome. Nodes with stronger connections receive more heat, facilitating a non-uniform yet structured spread of neural activity. The spectral decomposition of the diffusive laplacian can be seen as the building blocks mediating the diffusion of the neural activity (Atasoy et al., 2016). In contrast, the connectome harmonics of the random walk laplacian describe the probabilistic state for a given node to transfer information to another node. Thus, the connectome harmonics derived from the random walk Laplacian describe probabilistic transitions of neural activity between nodes. In this work, we considered the Laplacian (symmetric normalized laplacian, ie Diffusive) with the adjacency matrix weighted by strength of the structural connections. In the following discussion of the results, we interchangeably use the terms ”weaker coupling” and ”decoupling” when referring to the results of this study.

### 4.1 EEG in sensory areas is coupled to structural connectivity in movie-watching and resting state

A limited number of studies have undertaken the joint analysis of EEG cortical activity with structural connectomes using GSP, either with visual event-related potentials (Glomb et al., 2020; Rué-Queralt et al., 2021), auditory steady-state responses (Rué-Queralt et al., 2023) or during interictal epileptic discharges (Rigoni et al., 2023). Because the focus in these studies was on event-related experiments, the probed time windows were brief, ranging from a few hundred milliseconds to a few seconds. By contrast, our study investigates structure-function coupling with several minutes of continuous EEG and individual structural connectomes. Our findings (Fig 2c-d, Fig 6) indicate that the raw activity in visual, motor and auditory systems are coupled to the underlying structure across Video 1, 2, and Rest. These observations confirm and extend previous research (Glomb et al., 2020; Rué-Queralt et al., 2021, 2023), which has evaluated the contribution of structural connectivity eigenmodes in expressing the cortical EEG during visual and auditory tasks. Their observation that the sparse set of eigenmodes suffices to reconstruct the source activity implies that the low-frequency eigenmodes carry more signal power (i.e., more expressive) relative to their high-frequency counterpart. By definition of the SDI metric, this refers to coupling. Studies such as Glomb et al., 2020; Rué-Queralt et al., 2021 found that during visual-evoked tasks, the signal becomes more compact, i.e., the first few eigenmodes capture most of the signal power. In particular, Rué-Queralt et al., 2021, 2023 found that during visual and auditory tasks, the contributions of the low-frequency eigenmodes become dominant, also indicating coupling during the said tasks. As a consequence, our findings in sensory regions during continuous, long EEG segments complement the observations during isolated, shorter events, suggesting a consistent behavior of these systems during short and long time periods. In addition, existing work has estimated SDI with MEG during resting state, and also found that visual regions (among others) were strongly coupled to the structural connectome (Griffa & Preti, 2022).

From the individual analysis with distinct continuous cognitive states such as video and rest, we observed a largely similar trend consistently across cognitive states. This prompted a question whether the coupling relationship reorganizes selectively depending on task. To this end, we analyzed the coupling relationship between video and rest through a pairwise comparison. Results (Fig 2e) indicated that the raw activity during the movie exhibits stronger coupling across the cortex. In particular, late visual areas such as V3, V6, and somatosensory cortex exhibit stronger coupling during the movie, a trend consistent across both videos. This observation goes inline with the prior studies (Glomb et al., 2020; Rué-Queralt et al., 2021) in terms of more coupling during visual-evoked events relative to moments prior. Taken together, while the coupling relationship of the sensory systems is largely stable across cognitive states, the strength of the structure-function in selected systems appears to be modulated by the ongoing task condition.

### 4.2 Decoupling of EEG activity in parahippocampal and inferior frontal areas

Our finding that the parahippocampal area and inferior frontal areas were decoupled with the structure, a trend consistent across videos and during rest, aligns with the literature. In a study from Rigoni and colleagues analyzing EEG during four second-long epochs of interictal epileptic spikes decomposed on network harmonics (i.e. eigenvectors) of a structural connectome, the authors report that at least three patients have significant decoupling of ventral, parahippocampal areas as well as inferior frontal areas (Rigoni et al., 2023). In addition, prior work using resting-state MEG on 84 subjects have found the same regions to be decoupled according to the SDI (Griffa & Preti, 2022). Taken together, these results suggest decoupling of similar brain areas from short periods of four seconds in Rigoni’s study (Rigoni et al., 2023), and up to a few minutes of resting state and video watching in our results and in Griffa and Preti’s (Griffa & Preti, 2022). This questions the potential presence of time-varying aspects of structure-function coupling.

### 4.3 Temporal stability of structure-function coupling during video watching

Cognitive functions arise from the interplay between the brain dynamics and the network architecture. The static anatomy giving rise to the repertoire of functional activity that unfolds over time is fundamentally dynamic. We assessed how the structure supports the function over time in a region-resolved manner. We found that sensorimotor regions exhibit a stable relationship with the anatomy, whereas the transmodal regions such as vmPFC tend to show dynamic temporal fluctuation. Our findings build upon the previous work that showed temporal hierarchy (see Huntenburg et al., 2018 for review). This concept complements the spatial hierarchy of functional processing (Margulies et al., 2016) (i.e. spatial receptive fields), extending to the temporal domain, elucidating how the brain processes the information that is originating from the external world to self-oriented thoughts. In particular, temporal receptive fields characterize the hierarchy of temporal processing (Hasson et al., 2008). Accumulation of information over timescales is shown to vary across brain regions, with the sensory regions rapidly processing instantaneous sensory inputs. In contrast, the higher order system accumulates the information over a longer timescale (Baldassano et al., 2017; Hasson et al., 2008). The reasons for unimodal systems processing the input signal at a faster pace may suggest the critical nature of surviving and encountering dangerous situations (Kiebel et al., 2008), while the slow encoding in the transmodal cortices may imply that its inputs are not life-threatening. Minimal temporal fluctuation in the tethering of sensory systems can be seen as a highly reliable system that consistently relies on the structure, without much flexibility. Evolutionarily, requiring a reliable system to sample the input signal that puts an agent’s survival at risk, could be justified. We can also see in terms of functional specialization, as unimodal systems are specialized and tied to one particular task, thus expecting these systems to interact with the anatomy in the same way would be appropriate. On the other hand, the structure-function coupling varying at a higher rate in the transmodal cortices are flexible systems, reconfiguring to meet the task demands at hand. Interestingly, we found that a transmodal region PCC is associated with minimal temporal variation in structure-function association. PCC is a core DMN region, with the understanding of its role remaining to be fully understood (Leech & Sharp, 2014). One of the theories of its role is that it is controlling the balance between internal and external attention (Mesulam, 1998). It can be thought of as a ”watchdog”, and perhaps essential to be stable temporally to make the transition. Future studies are necessary to confirm or refute this finding.

### 4.4 Similarity of structure-function coupling across EEG frequency bands

We also analyzed whether the SDI patterns differ across distinct EEG frequency bands (Fig 4, first four columns). Prior research on MEG data at rest (Griffa & Preti, 2022) observed a high spatial topography across *δ* to *β* bands, supporting our observation of strong correlation in the SDI values between frequency bands. This suggests consistent structure-function association across frequency bands. Comparing between conditions, regions such as the motor cortices and Retrosplenial complex in the PCC exhibited stronger coupling to the structure during the movie across all frequency bands (Fig 4 last column). In Rué-Queralt et al., 2023, the authors defined a joint transform over graph and frequency domain by combining structural harmonics with Morlet wavelets (called ”time vertex spectral representation” in their paper), and applied to EEG in visual and auditory tasks. They found that slower EEG rhythms were mostly coupled to the structure, while the activity of faster temporal frequencies tended to be more decoupled (Rué-Queralt et al., 2023). In contrast, we found regions such as parahippocampus, inferior frontal junction to be consistently decoupled across slow and fast EEG rhythms during several minutes of EEG. This difference might arise from the variation in window duration, which was around a few seconds in Rué-Queralt et al., 2023, compared to substantially longer durations in our study and in Griffa and Preti’s study (Griffa & Preti, 2022). One might wonder if applying Hilbert Transform obscures the subtle oscillations that cause this similarity. We investigated this by considering the bandpass signal, and fourier coefficients computed by STFT (see supplementary material, section11). As with the envelope signal, we observed similar spatial topography across frequency bands. These results suggest that the anatomy constrains the slower and faster rhythms in a similar way. These findings are supported when analyzed with a) envelope signal, b) bandpass signals, and c) STFT coefficients, and these findings are inline also with the previous studies (Griffa & Preti, 2022).

### 4.5 Spatially focal coupling relationship in higher-order systems compared to haemodynamics

Here we relate the coupling relationship of the EEG SDI results with the existing work analyzing haemodynamic responses, to assess the possibility of a similar structure-function coupling at two distinct temporal scales. The question of whether the coupling relationship of the brain response at a slower time scale (e.g: haemodynamic) and the electrophysiological response exhibit an overlapping trend has not been fully studied before. One study from Griffa and Preti (Griffa & Preti, 2022) with MEG data showed that the group-averaged SDI patterns deviated from studies with fMRI (Preti & Van De Ville, 2019). Numerous studies have shown and confirmed the global hierarchy from unimodal-transmodal in different contexts, ranging from cortical microstructure (Huntenburg et al., 2017), to macroscale connectivity (Bernhardt et al., 2022; Margulies et al., 2016) and structure-function coupling (Preti & Van De Ville, 2019)(see Huntenburg et al., 2018 for a review). Our findings (Fig 6) show that the cortical activity in the sensorimotor areas during both videos and rest is generally coupled to the structure, supporting prior research. In particular, studies such as (Margulies et al., 2016; Preti & Van De Ville, 2019; Yang et al., 2023) analyzed the macroscale organization or the coupling relationship using resting-state fMRI, and generally found a gradient-like spatial organisation from sensorimotor systems to transmodal ones. Preti and Van de Ville (Preti & Van De Ville, 2019) observed that the three systems exhibiting strong coupling include Visual, Motor, and Auditory, and some of the systems exhibiting strong decoupling include Emotion, Social Cognition, and Memory. Situating these systems in the context of EEG reveals that they are very much in agreement not only at Rest but also during both videos, and a similar pattern was found in MEG (Griffa & Preti, 2022). However, there are two key differences in our functional decoding results (Fig 6). The first one is the relative place of the auditory system, which was less coupled than the motor and visual systems, in both Rest and Video, which suggests flexibility in the coupling of the auditory system with respect to the structure. Additionally, transmodal systems such as Social, Memory-retrieval, and Emotion were associated in our EEG results with a wider range of structure-function patterns ranging from strongly coupled to weakly coupled areas. Taken together, our results with EEG confirm previous observations with fMRI in the sensorimotor cortex, while it contributes to understanding more about the dynamics of EEG in the transmodal cortices, with the spatially refined structure-function relationship.

### 4.6 The topography of EEG structure-function coupling is consistent across movie-watching and resting-state

Movies can elicit a richer repertoire of brain states than resting-state, providing a window to understand both exteroceptive and interoceptive processes (Meer et al., 2020). Whether the cortical organization during the movie differs from the resting state has been previously studied (Kringelbach et al., 2023; Samara et al., 2023). For example, Samara et al. 2023 show that cortical organization during movie-watching follows a different principle than during rest, with the gradients being modality-specific. They found that during movie-watching the default mode network and frontoparietal network formed a heteromodal peak, while the sensorimotor regions anchored the unimodal pole for the first three fMRI gradients. Another fMRI study from Kringelbach and colleagues (Kringelbach et al., 2023) investigated the hierachical reorganisation of the functional brain activity during movie compared to rest, using a generative model of effective connectivity. This approach enables to model the direction of information flow, and estimate the hierarchy of activity as a change of balance in causal interactions between brain regions, under different task conditions of movies and rest. Their findings indicate that fMRI activity is relatively less hierarchical during movie than during rest, as they beautifully put it as ”flattening of the hierarchy”. This sparks the question as to whether the coupling relationship is also reshaped between movie and rest. While there is no study directly comparing structure-function coupling in fMRI using both resting-state and video watching, our findings using EEG suggest a similar spatial distribution of coupling across tasks. Specifically, sensory systems strongly coupled to the structure are apparent across all contexts, while the coupling relationship in the higher-order systems is disrupted (i.e. spans between coupling and decoupling, Fig 6). When performing a pairwise comparison between movie and rest however, we found the coupling relationship was strengthened watching a movie.These findings were reliable when tested with Video 2, as indicated by the ICC score for the contrast condition. However, upon closer look we can observe hemispheric asymmetries contrasting Video 1 and Video 2 with the rest. We found that the regions in the left anterior temporal lobe were exhibiting strengthened coupling while watching videos (1 and 2) relative to rest, which, in the other hemisphere, were not observed to be significantly different from the null models. One might wonder why, and perhaps lean to attributing it to functional specialization in the specific hemisphere. However, given that the EEG signal is less known for its spatial resolution, interpreting the behavior of the isolated region requires more validation. Future studies using better imaging techniques, and other source localization techniques may clarify how the structure supports a certain task relative to the resting state.

### 4.7 Structure-Function coupling across cognitive and conscious states

Our findings from the normal wakeful conscious state complement the previous knowledge from altered states of consciousness (ASC), providing a broader picture to understand the brain dynamics. Connectome harmonics have the potential to characterize structure-function association across cognitive states, as in our study. Besides, it has been extended to characterizing how structural architecture shapes function across different conscious states, including ASC such as those induced by anesthesia, ketamine, and psychedelics (see Luppi et al., 2024 for review). Broadly, these states are characterized by a prevalence of low-frequency harmonics for the Anesthetic state, and high-frequency harmonics for Psychedelics-induced states. In particular, elevated energy in high-frequency harmonics is observed in altered states induced by psychoactive substances such as Ketamine (Luppi et al., 2023), LSD (Atasoy et al., 2017), Psilocybin (Atasoy et al., 2018). In contrast, the opposite effect is seen in altered states induced by compounds such as Anesthesia under Propofol (Luppi et al., 2023). These studies highlight the dominant role of low to high-frequency harmonics across the Anesthesia to Psychedelics states. Normal wakeful state has to sit somewhere along this hierarchy. Previous studies characterized the significant contribution of connectome harmonics primarily using graph power or graph energy. Our results comparing normal resting and movie-watching states directly provide insights on this front. We found no significant difference in the graph Power spectrum between these two states. This indicates that, under normal conscious state, the contribution of connectome harmonics is similar between movie-watching and resting state. However, from Mediano, Rosas, and colleagues analyzing the effect of external stimulation on the psychedelic experience indexed by Lempel-Ziv complexity, we can observe noticeable differences between the same cognitive states (Mediano et al., 2024). This could suggest a fundamental distinction between normal and altered states of consciousness, with the brain operating in a controlled manner without significant differences across cognitive states, as opposed to altered states where the cognition is less meticulous and more disordered (Carhart-Harris et al., 2014).

### 4.8 Limitations and perspectives

Several limitations need to be acknowledged, as they may be useful to consider in follow-up research building on our work. An important limitation to consider is how source-estimation of the EEG influences the graph power spectrum. Inverse methods impose constraints that result in a spatially smooth source time course. This can lead to concentration of the power mostly captured by the lower end of the connectome harmonics. This pattern is consistent with findings from several other EEG studies employing different families of inverse methods, therefore the lower Eigenmode dominance does not fully account for source reconstruction and its methods. Hence, we suggest interpreting our findings in the light of this factor. Secondly, we have used a template for the spatial definition of ROIs, using the HCP-MMP atlas (Glasser et al., 2016), known to have a good generalization across subjects. However, results might differ when using individually defined parcels or hyper-alignment methods (Xu et al., 2012; Zhou et al., 2022). This aspect could be the subject of future work, in particular with datasets that are adapted to estimate individual parcellations and connectomes (Pascucci, Tourbier, et al., 2022). Finally, we were unable to show clear differences in the topography of the SDI pattern as a function of classical EEG frequency bands. An interesting perspective could revisit this result by reparameterizing the EEG power spectrum using aperiodic and periodic components (Donoghue et al., 2020).

### 4.9 Conclusions

In this study, we provide a comprehensive characterization of the structure-function relationship based on diffusion imaging and cortical EEG recorded while participants either rested or watched a movie. We quantified the coupling patterns using subject-specific Eigenmodes of individual structural connectomes using graph signal processing. Our findings suggest a similar spatial topography of coupling of sensory systems to the structure across video watching and rest, complementing prior results obtained with fMRI. We also found a few regions to be consistently decoupled from the structure, namely the parahippocampal cortex and superior parietal areas, and an overall strengthening of coupling in movie watching compared to resting-state. Our results obtained with continuous EEG (several minutes of movie-watching and rest) also bring complementary insights with respect to previous work that has analyzed the link between EEG and structural connectivity during event-related short data segments. Using functional decoding we showed that the cortical activity in the unimodal systems were associated to strong coupling, inline with previous fMRI findings, whereas the higher-order systems such as memory retrieval and emotions were associated between coupling and decoupling, bringing in novel insights compared to previous fMRI work.

## 5 CRediT Statement

**Venkatesh Subramani**: Conceptualization, Methodology, Software, Validation, Formal Analysis, Investigation, Writing - Original Draft, Writing - Review & Editing, Visualization. **Giulia Lioi**: Conceptualization, Methodology, Resources, Writing - Review & Editing, Supervision, Project administration. **Karim Jerbi**: Conceptualization, Writing - Review & Editing, Supervision, Project administration. **Nicolas Farrugia**: Conceptualization, Methodology, Resources, Writing - Review & Editing, Supervision, Project administration.

## 7 Ethics Statement

The data used in this study is collected by HBN, which obtained written consents from the all the participants to openly share the data following a de-identification process.

## 8 Code and Data availability

All the scripts necessary to perform analysis are shared on Github

## 9 Acknowledgements

EEG and DWI data were shared as part of the Healthy Brain Network. We thank Maxmillian Nentwich for providing us the list of subjects having good quality EEG identified in their study (Nentwich et al., 2020). We are grateful to Maria-Giulia Preti for sharing us the details about the pipeline utilized in their study (Preti & Van De Ville, 2019) for constructing structural connectome. Our Qsiprep pipeline was run in a High-Performance Cluster grid thanks to the BC DRI Group, Calcul québec and the Digital Research Alliance of Canada (https://alliancecan.ca/) and we appreciate their continued support and swift action in fixing a problem during the analysis. We also thank Alix Lamouroux for helping us get started with Qsiprep, and Yassine El Ouahidi for providing us the script to perform decoding with NiMare.

K.J. is supported by funding from the Canada Research Chairs (950-232368) program and a Discovery Grant from the Natural Sciences and Engineering Research Council of Canada (2021-03426).

## Supplementary material

### 10 Alternative strategy for computing cut-off frequency adapted to EEG demonstrates tangible way to compute SDI

The cut-off frequency that characterizes low-frequency and high-frequency Eigenmodes was C = 2 across task conditions. Lower Eigenmodes carrying high power may be attributed to the spatial smoothness introduced by the source localization algorithm. Previous studies such as (Preti & Van De Ville, 2019) analyzing the structure-function association of fMRI signals preserved the smoothest Eigenmode to define the cut-off frequency. Previous EEG studies using different family of source localization algorithms revealed that the first connectome harmonics expresses the cortical activity disproportionately (Glomb et al., 2020; Rué-Queralt et al., 2021).

Thus, we re-evaluated the choice of the cut-off by considering a different criterion, that only considers the graph spectrum for the non-constant eigenvectors.

We computed the SDI for the raw EEG signal following this strategy. The group-level Structural-Decoupling Index (SDI) results for the raw EEG signal (1-62.5 Hz) are presented in Fig 7. The median cut-off is shifted, C = 14 for Video 1, C = 13 for Rest, and C = 14 for Video 2 (compared to C = 2 earlier). A detailed breakdown of the new results and the comparison with the results in the manuscript are stated below.

**Table 1:** Differences in Coupling as a function of Task conditions and cut-off frequencies. (*C* = 2: Manuscript)

| **Video 1** |  | **Rest** |  | **Video 2** |  |
| --- | --- | --- | --- | --- | --- |
| $C = 14$ | $C = 2$ | $C = 13$ | $C = 2$ | $C = 14$ | $C = 2$ |
| Primary somatosensory cortex (Brodmann Area 1, SDI = -2.24) | Primary Somatosensory cortex (Area 2; SDI: -2.24) | Primary Motor Cortex (SDI = -2.20) | Primary somatosensory cortex (BA1, SDI: -1.78) | Primary somatosensory cortex (Area 2; SDI: -2.2) | Primary somatosensory cortex (Brodmann Area 1, SDI = -1.79) |
| Ventral anterior cingulate (SDI = -2.22) | Primary Sensory cortex (SDI: -1.8) | Primary somatosensory cortex (Brodmann Area 1, SDI = -2.24) | Visual V2 (SDI: -1.74) | Ventral anterior cingulate (SDI: -2.22) | Visual V3 (SDI: -1.77) |
| Brodmann area 47 in the frontal region SDI = -2.18 | Second Visual area (V2; SDI: -1.8) | Third Visual Area (SDI: -2.12) | Visual V3 (SDI: -1.74) | Primary Motor Cortex (SDI: -2.18) | Visual V2 (SDI: -1.77) |

**Figure 7:**
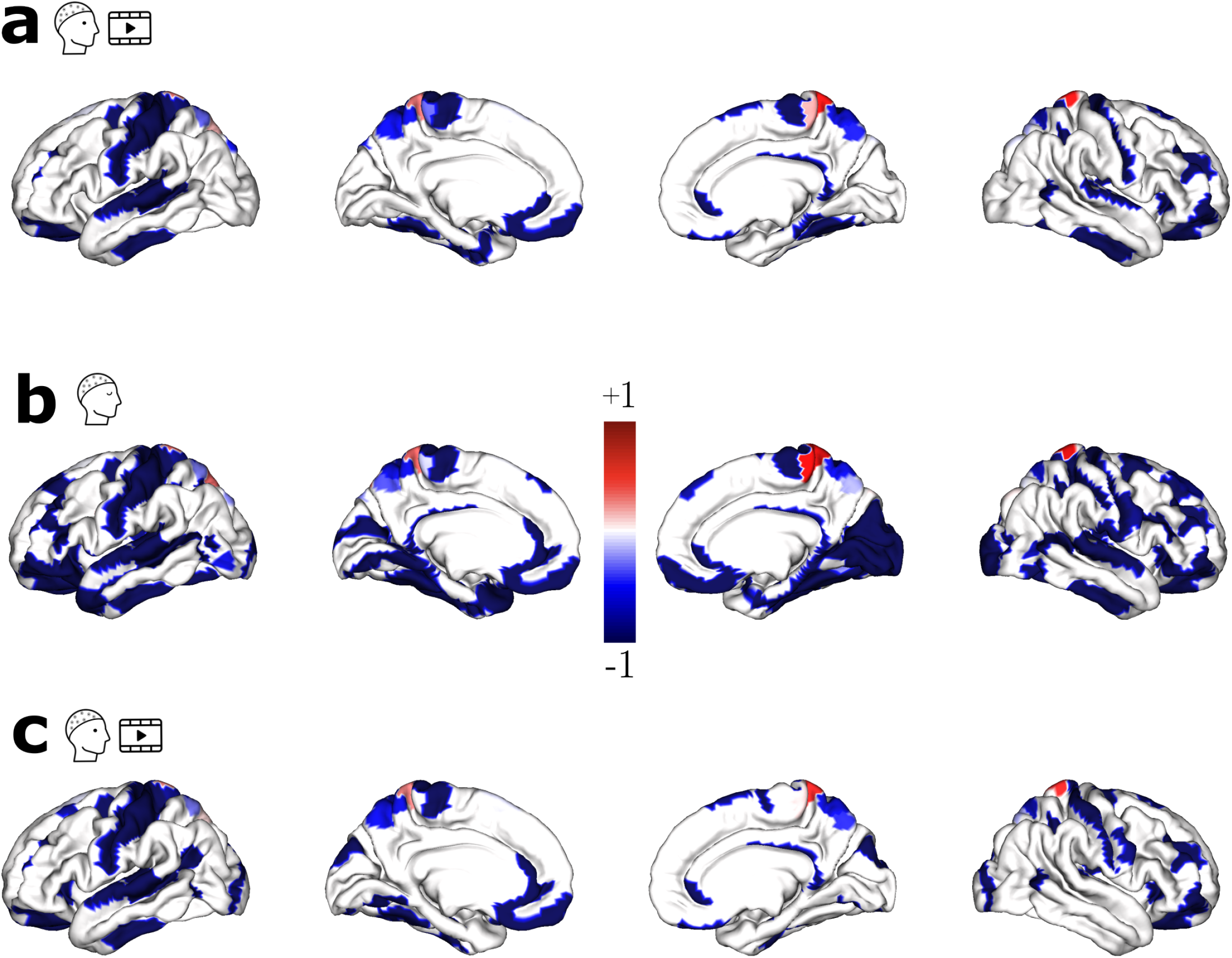
Group level SDI maps with the cut-off frequency chosen with a different criterion (i.e. disregarding the first constant eigenvector and choosing the frequency that dichotomizes the power spectrum of non-constant eigenvectors) (**a** Video1, **b** Rest, **c** Video2)

**Table 2:** Differences in Decoupling as a function of Task conditions and cut-off frequencies. (*C* = 2: Manuscript)

| Video 1 |  | Rest |  | Video 2 |  |
| --- | --- | --- | --- | --- | --- |
| $C = 14$ | $C = 2$ | $C = 13$ | $C = 12$ | $C = 14$ | $C = 2$ |
| Parahippocampal area 2 (SDI = 0.64) | Parahippocampal Area 2 (SDI: 0.62) | Parahippocampal area 2 (SDI = 0.61) | Parahippocampal area 2 (SDI: 0.71) | Posterior parietal cortex (SDI: 0.39) | Inferior Frontal Sulcal area (IFJp; SDI: 0.62) |
| Posterior parietal cortex (Area 5L; SDI: 0.38) | Inferior Frontal Sulcal area (IFJp; SDI: 0.5) | Posterior parietal cortex (Area 5L; SDI: 0.5) | Inferior Frontal Sulcal area (IFJp; SDI: 0.6) | Area 5m of Superior parietal lobule (SDI: 0.06) | Parahippocampal Area 2 (SDI: 0.49) |
| Area 5m of Superior parietal lobule (SDI: 0.17) | Posterior parietal cortex (Area 5L; SDI: 0.2) | Area 5m of Superior parietal lobule (SDI: 0.49) | Subgenual Anterior Cingulate cortex (Area s32; SDI: 0.38) | Lateral Area 7P of Superior parietal lobule (SDI: 0.07) | Subgenual Anterior Cingulate cortex (Area s32; SDI: 0.26) |

**Table 3:** Contrast map between task conditions using different cut-off frequencies (*C* = 2: Manuscript)

| Video 1 vs Rest |  | Video 2 vs Rest |  |
| --- | --- | --- | --- |
| $C = 14$ | $C = 2$ | $C = 14$ | $C = 2$ |
| Area 5m of Superior parietal lobule ($t(42) = -4.65$) | Visual area 7 (V7; $t(42) = -5$) | Visual area V6A ($t(42) = -5.5$) | Visual areas (V3A, V6A; $t(42) = -5.4, -5.4$ respectively) |
| Visual area V6A ($t(42) = -4.5$) | Inferior parietal cortex (Area PGP; $t(42) = -4.65$) | Area 5m of Superior parietal lobule ($t(42) = -4.5$) | Area 5m of Superior parietal lobule ($t(42) = -5.2$) |

**Figure 8:**
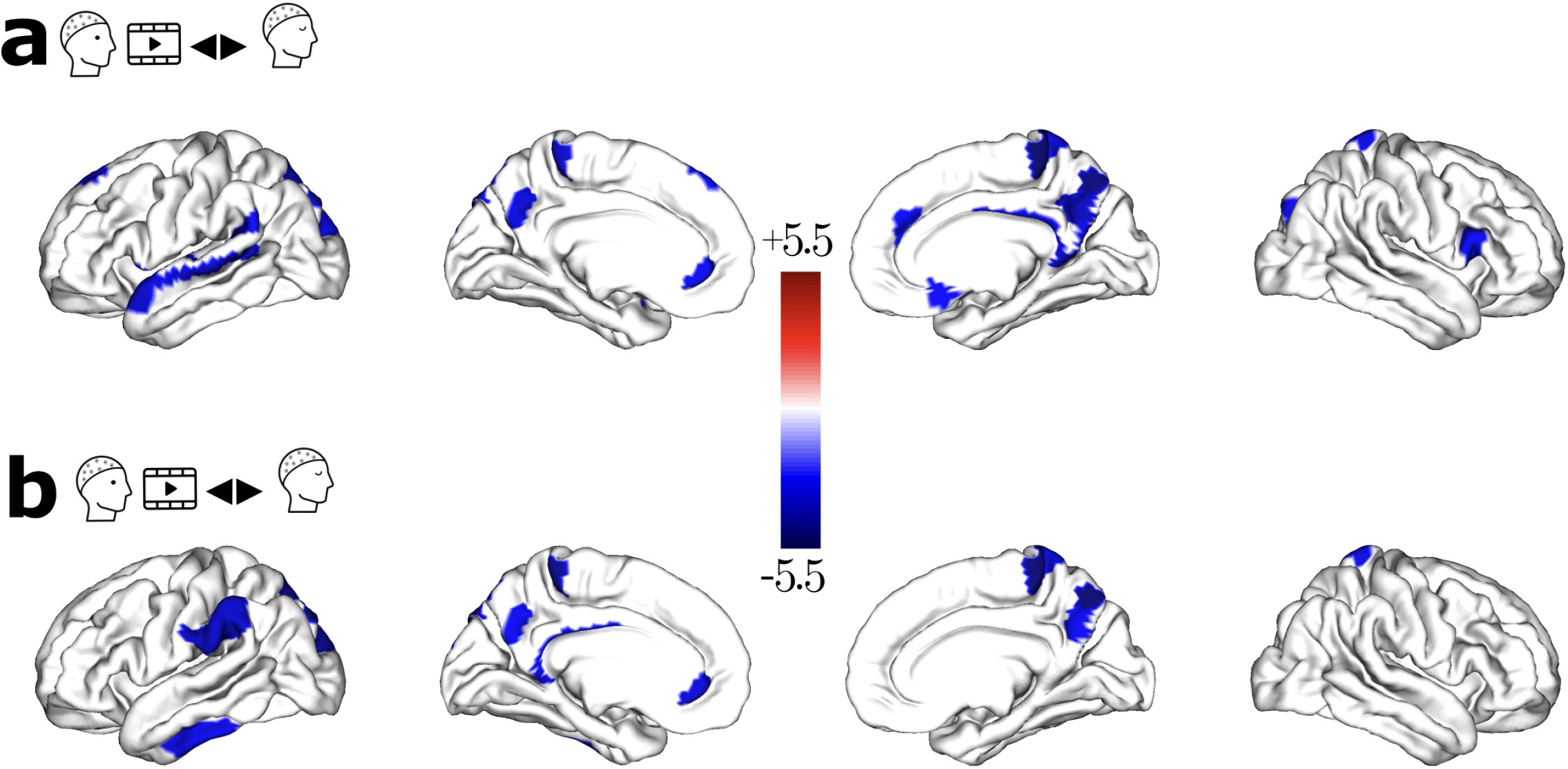
Contrast map between task conditions using the new cutoff strategy (**a** Video1 vs Rest, **b** Video2 vs Rest)

Overall, the revised criterion has resulted in changes in SDI results, particularly in the cut-off values and spatial patterns of coupling and decoupling across different brain regions. Then, we sought to see the coherence of adapting the SDI computation for EEG by performing Neurosynth/Nimare decoding as well (Fig 9). These results can be interpreted as follows: sensory systems are associated with coupling, while the higher-order systems tend to run across the coupling-decoupling gradient. Albeit differences (higher order systems have high Zscore on the decoupling bin, as opposed to rather uniform spread out), our original interpretation still holds true.

**Figure 9:**
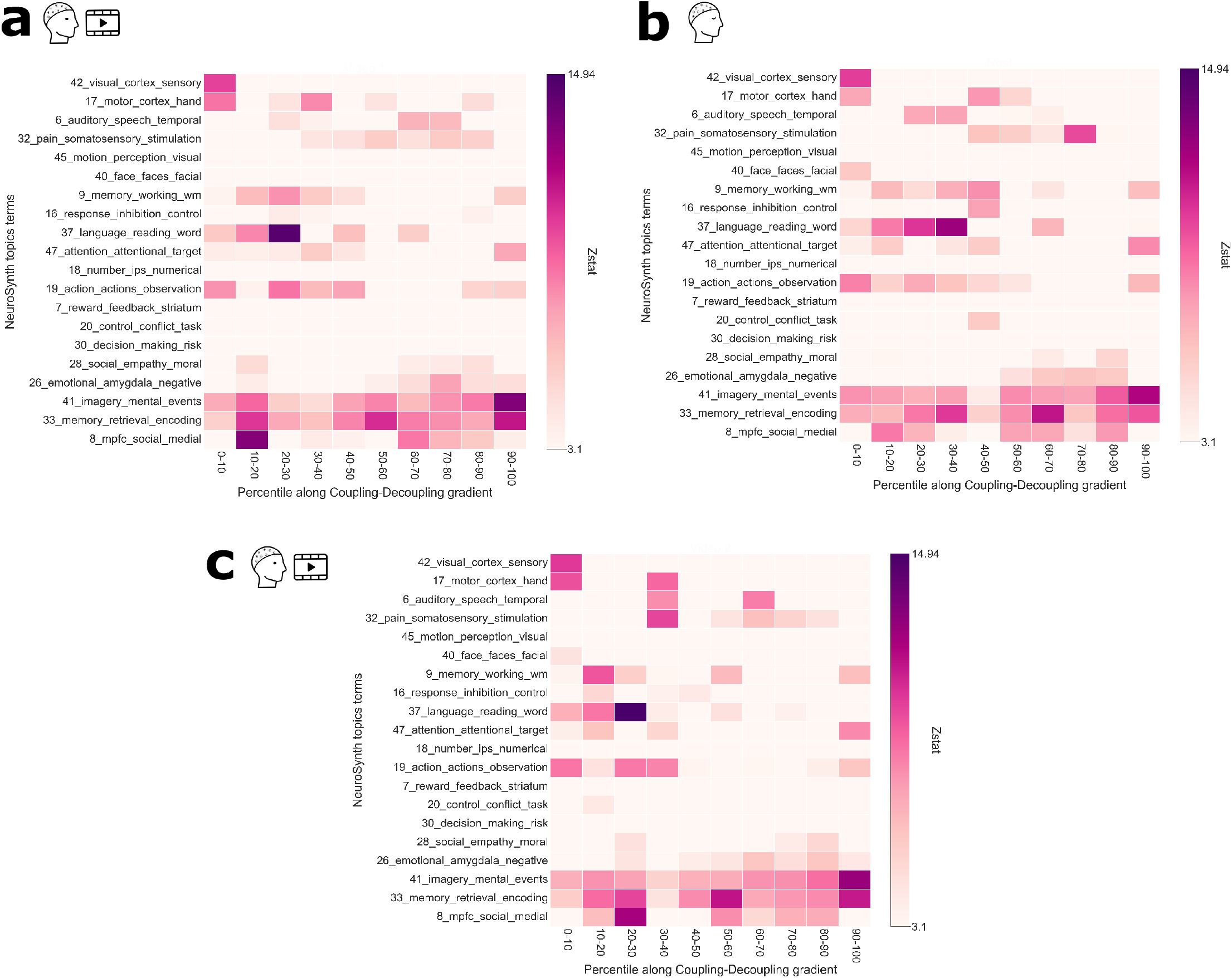
Decoding the spatial maps for different task conditions (**a** Video 1, **b** Rest, **c** Video 2) for which the SDI cutoff frequency is computed, excluding the constant eigenvector.

These results after exclusion of the power of the constant eigenvector, and considering the whole power spectrum in the manuscript being complementary speak to the robustness of the methodology and the findings. The new setup could demonstrate a tangible way to approach the source-localized EEG signals.

## 11 Structure-Function coupling quantified for bandpass signals and Fourier coefficients reveals similar spatial distribution across frequency bands

We observed similar topography across frequency bands when the structure-function coupling is quantified for the envelope signal. One might wonder if applying Hilbert Transform obscures the subtle oscillations that cause this similarity. We investigated this by considering the bandpass signal, and fourier coefficients computed by Short-Term Fourier Transform (STFT).

We computed the SDI for the bandpass signal, and performed statistical thresholding with the same pipeline. Group-level maps are presented in Fig 10 for the various EEG frequency bands ranging from *θ* (4 - 8) to *γ* (30 40 Hz). We compared the spatial distribution across all pairs of frequency bands using group-level maps. Results reveal a high correlation with the average Spearman’s rho of 0.84 across all pairs (SD: 0.02). This is comparable to the envelope signal (average rho being 0.86, SD being 0.02).

**Figure 10:**
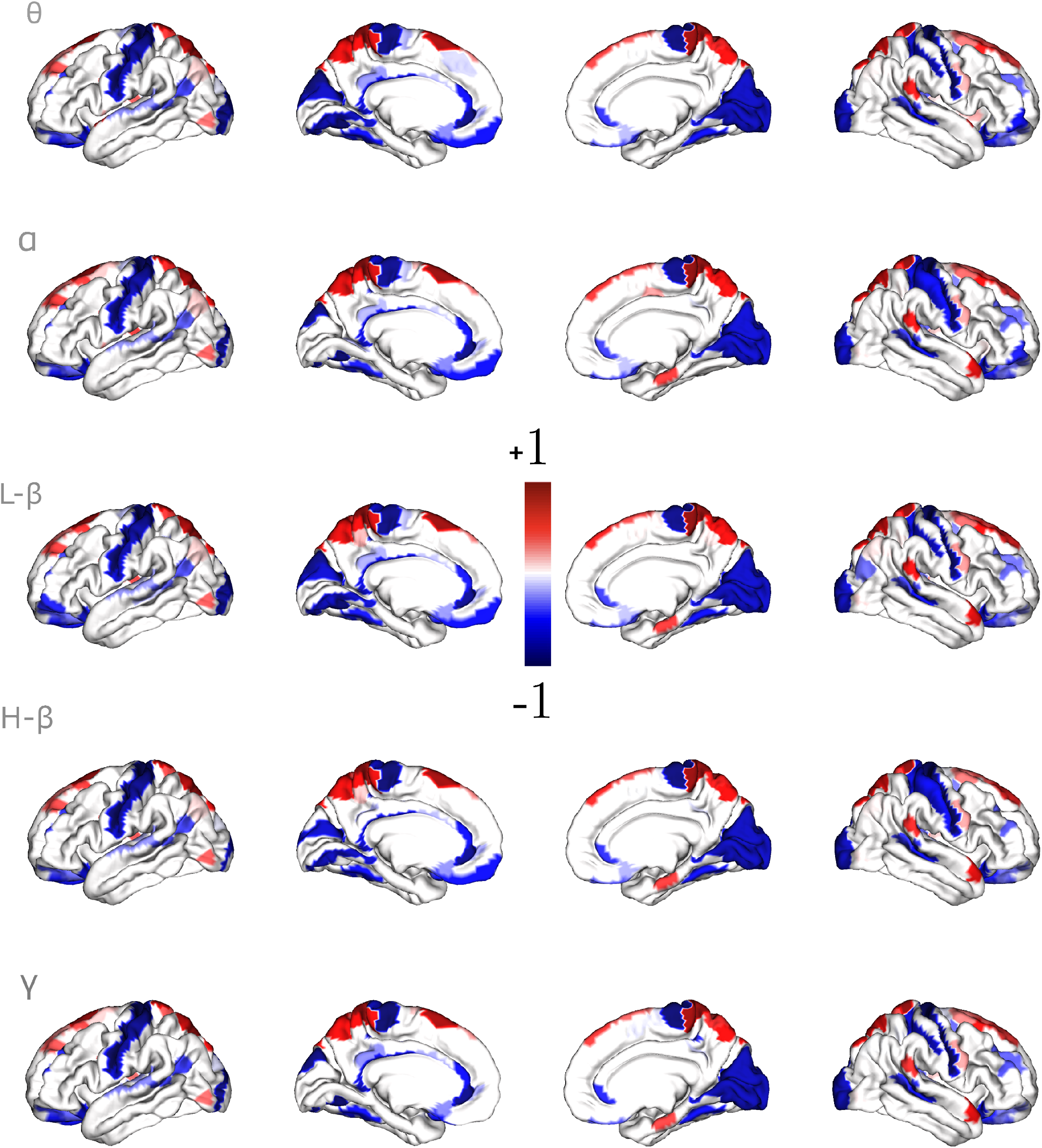
Group SDI maps of the link between bandpass filtered signals and the anatomy

We analyzed the time and frequency characteristics of a source-localized EEG signal using the Short-time Fourier Transform (STFT). Specifically, we transformed the EEG data into time-frequency coefficients using a window length of 200 ms, a 50% overlap, and Hann windows. This window length was chosen as it offers a reasonable balance between the scale of frequencies (at least 5Hz) and the time samples (11 per second). These coefficients were then considered as the graph signal, and the structure-function coupling was quantified. When necessary, coefficients were averaged within the frequency range (e.g., two coefficients in the high-*β* range between 20 and 30 Hz). Group-level maps are presented in Fig 11 for the various EEG frequency bands ranging from *θ* (4 - 8) to *γ* (30 - 40 Hz). We computed the whole-brain spatial similarity across all pairs of frequency bands at the group level. Results reveal the average rho being 0.73 (SD: 0.13). Though there is a slight drop in averaged rho, the spatial similarity is nevertheless high.

**Figure 11:**
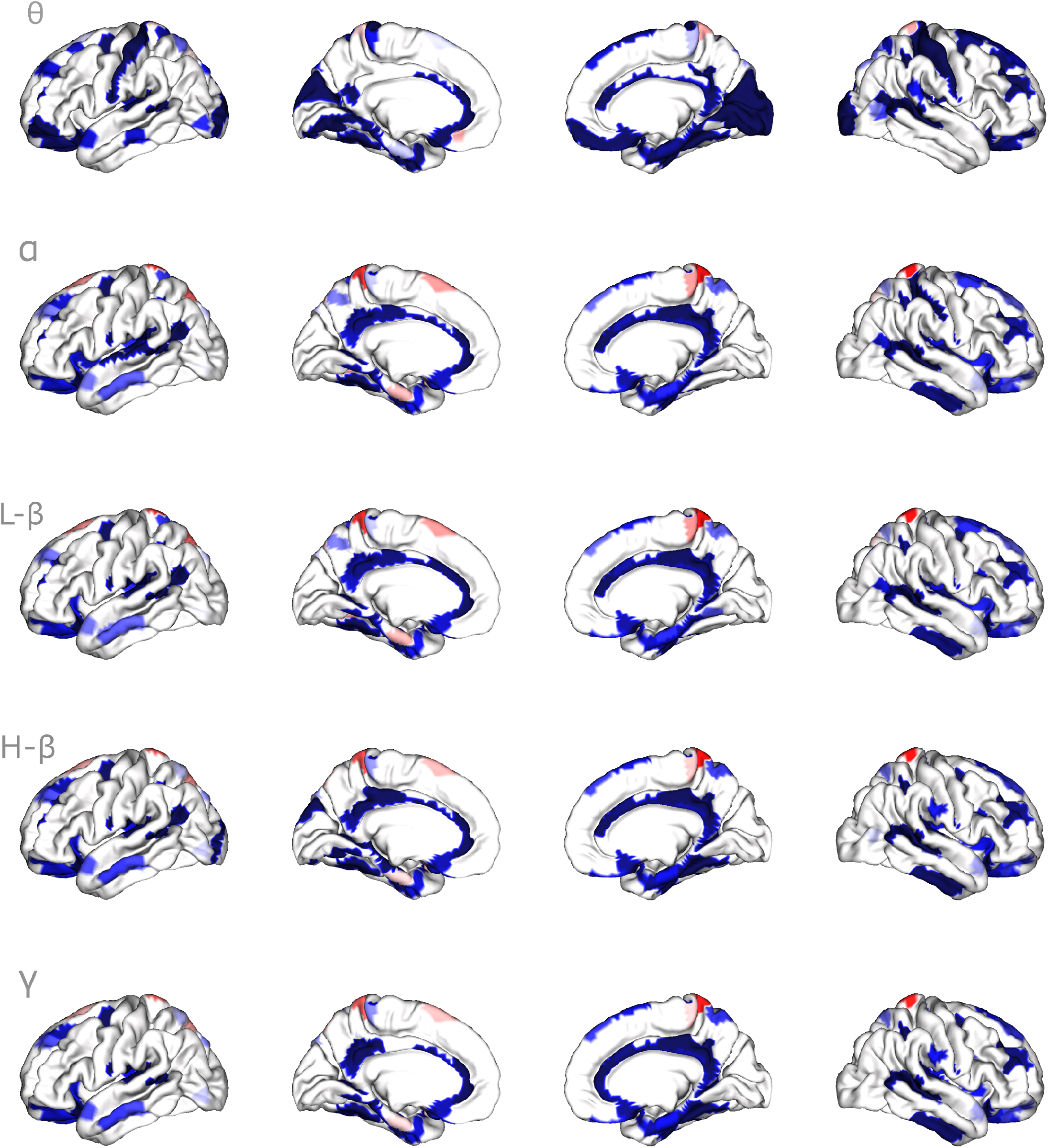
Group level SDI maps depicting the link between the STFT coefficients of the cortical signal and the anatomy.

Despite considering the bandpass signal, and STFT coefficients as graph signals, we still find comparable spatial similarity, indicating the oscillation that might be preserved is not sufficient for the SDI metric to show clear differences.

## Footnotes

1 https://www.youtube.com/watch?v=HNXxJIhVALI

2 https://www.youtube.com/watch?v=XwWyTts06tU

## References

Atasoy, S., Donnelly, I., & Pearson, J. (2016). Human brain networks function in connectome-specific harmonic waves. Nature communications, 7 (1), 10340.

Atasoy, S., Roseman, L., Kaelen, M., Kringelbach, M. L., Deco, G., & Carhart-Harris, R. L. (2017). Connectome-harmonic decomposition of human brain activity reveals dynamical repertoire re-organization under lsd. Scientific reports, 7 (1), 17661.

Atasoy, S., Vohryzek, J., Deco, G., Carhart-Harris, R. L., & Kringelbach, M. L. (2018). Common neural signatures of psychedelics: Frequency-specific energy changes and repertoire expansion revealed using connectome-harmonic decomposition. Progress in Brain Research, 242, 97–120.

Bajada, C. J., Costa Campos, L. Q., Caspers, S., Muscat, R., Parker, G. J., Lambon Ralph, M. A., Cloutman, L. L., & Trujillo-Barreto, N. J. (2020). A tutorial and tool for exploring feature similarity gradients with MRI data. NeuroImage, 221 (January), 117140. 10.1016/j.neuroimage.2020.117140

Baldassano, C., Chen, J., Zadbood, A., Pillow, J. W., Hasson, U., & Norman, K. A. (2017). Discovering event structure in continuous narrative perception and memory. Neuron, 95 (3), 709–721.

Bassett, D. S., & Sporns, O. (2017). Network neuroscience. Nature neuroscience, 20 (3), 353–364.

Belkin, M., & Niyogi, P. (2003). Laplacian eigenmaps for dimensionality reduction and data representation. Neural computation, 15 (6), 1373–1396.

Bernhardt, B. C., Smallwood, J., Keilholz, S., & Margulies, D. S. (2022). Gradients in brain organization. NeuroImage, 251, 118987.

Carhart-Harris, R. L., Leech, R., Hellyer, P. J., Shanahan, M., Feilding, A., Tagliazucchi, E., Chialvo, D. R., & Nutt, D. (2014). The entropic brain: A theory of conscious states informed by neuroimaging research with psychedelic drugs. Frontiers in human neuroscience, 8, 55875.

Cieslak, M., Cook, P. A., He, X., Yeh, F.-C., Dhollander, T., Adebimpe, A., Aguirre, G. K., Bassett, D. S., Betzel, R. F., Bourque, J., et al. (2021). Qsiprep: An integrative platform for preprocessing and reconstructing diffusion mri data. Nature methods, 18 (7), 775–778.

Dhollander, T., Mito, R., Raffelt, D., & Connelly, A. (2019). Improved white matter response function estimation for 3-tissue constrained spherical deconvolution. Proc. Intl. Soc. Mag. Reson. Med, 555 (10).

Donoghue, T., Haller, M., Peterson, E. J., Varma, P., Sebastian, P., Gao, R., Noto, T., Lara, A. H., Wallis, J. D., Knight, R. T., et al. (2020). Parameterizing neural power spectra into periodic and aperiodic components. Nature neuroscience, 23 (12), 1655–1665.

Faskowitz, J., Moyer, D., Handwerker, D. A., Gonzalez-Castillo, J., Bandettini, P. A., Jbabdi, S., & Betzel, R. (2023). Commentary on pang et al. (2023)nature. 10.1101/2023.07.20.549785

Fischl, B. (2012). Freesurfer. Neuroimage, 62 (2), 774–781.

Fotiadis, P., Parkes, L., Davis, K. A., Satterthwaite, T. D., Shinohara, R. T., & Bassett, D. S. (2024). Structure–function coupling in macroscale human brain networks. Nature Reviews Neuroscience, 1–17.

Glasser, M. F., Coalson, T. S., Robinson, E. C., Hacker, C. D., Harwell, J., Yacoub, E., Ugurbil, K., Andersson, J., Beckmann, C. F., Jenkinson, M., et al. (2016). A multi-modal parcellation of human cerebral cortex. Nature, 536 (7615), 171–178.

Glomb, K., Kringelbach, M. L., Deco, G., Hagmann, P., Pearson, J., & Atasoy, S. (2021). Functional harmonics reveal multi-dimensional basis functions underlying cortical organization. Cell Reports, 36 (8).

Glomb, K., Queralt, J. R., Pascucci, D., Defferrard, M., Tourbier, S., Carboni, M., Rubega, M., Vulliemoz, S., Plomp, G., & Hagmann, P. (2020). Connectome spectral analysis to track eeg task dynamics on a subsecond scale. NeuroImage, 221, 117137.

Gramfort, A., Luessi, M., Larson, E., Engemann, D. A., Strohmeier, D., Brodbeck, C., Goj, R., Jas, M., Brooks, T., Parkkonen, L., & Hämäläinen, M. S. (2013). MEG and EEG data analysis with MNE-Python. Frontiers in Neuroscience, 7 (267), 1–13. 10.3389/fnins.2013.00267

Griffa, A., & Preti, M. G. (2022). Brain structure-function coupling is unique to individuals across multiple frequency bands: A graph signal processing study. 2022 30th European Signal Processing Conference (EUSIPCO), 942–946.

Hagmann, P., Cammoun, L., Gigandet, X., Meuli, R., Honey, C. J., Wedeen, V. J., & Sporns, O. (2008). Mapping the structural core of human cerebral cortex. PLoS biology, 6 (7), e159.

Hasson, U., Yang, E., Vallines, I., Heeger, D. J., & Rubin, N. (2008). A hierarchy of temporal receptive windows in human cortex. Journal of neuroscience, 28 (10), 2539–2550.

Honey, C. J., Sporns, O., Cammoun, L., Gigandet, X., Thiran, J.-P., Meuli, R., & Hagmann, P. (2009). Predicting human resting-state functional connectivity from structural connectivity. Proceedings of the National Academy of Sciences, 106 (6), 2035–2040.

Huntenburg, J. M., Bazin, P.-L., Goulas, A., Tardif, C. L., Villringer, A., & Margulies, D. S. (2017). A systematic relationship between functional connectivity and intracortical myelin in the human cerebral cortex. Cerebral Cortex, 27 (2), 981–997.

Huntenburg, J. M., Bazin, P.-L., & Margulies, D. S. (2018). Large-scale gradients in human cortical organization. Trends in cognitive sciences, 22 (1), 21–31.

Kiebel, S. J., Daunizeau, J., & Friston, K. J. (2008). A hierarchy of time-scales and the brain. PLoS computational biology, 4 (11), e1000209.

Koo, T. K., & Li, M. Y. (2016). A guideline of selecting and reporting intraclass correlation coefficients for reliability research. Journal of chiropractic medicine, 15 (2), 155–163.

Kringelbach, M. L., Perl, Y. S., Tagliazucchi, E., & Deco, G. (2023). Toward naturalistic neuroscience: Mechanisms underlying the flattening of brain hierarchy in movie-watching compared to rest and task. Science Advances, 9 (2), eade6049.

Langer, N., Ho, E. J., Alexander, L. M., Xu, H. Y., Jozanovic, R. K., Henin, S., Petroni, A., Cohen, S., Marcelle, E. T., Parra, L. C., et al. (2017). A resource for assessing information processing in the developing brain using eeg and eye tracking. Scientific data, 4 (1), 1–20.

Leech, R., & Sharp, D. J. (2014). The role of the posterior cingulate cortex in cognition and disease. Brain, 137 (1), 12–32.

Lioi, G., Gripon, V., Brahim, A., Rousseau, F., & Farrugia, N. (2021). Gradients of connectivity as graph fourier bases of brain activity. Network Neuroscience, 5 (2), 322–336.

Luppi, A. I., Rosas, F. E., Mediano, P. A., Demertzi, A., Menon, D. K., & Stamatakis, E. A. (2024). Unravelling consciousness and brain function through the lens of time, space, and information. Trends in Neurosciences.

Luppi, A. I., Vohryzek, J., Kringelbach, M. L., Mediano, P. A., Craig, M. M., Adapa, R., Carhart-Harris, R. L., Roseman, L., Pappas, I., Peattie, A. R., et al. (2023). Distributed harmonic patterns of structure-function dependence orchestrate human consciousness. Communications biology, 6 (1), 117.

Lynn, C. W., & Bassett, D. S. (2019). The physics of brain network structure, function and control. Nature Reviews Physics, 1 (5), 318–332.

Margulies, D. S., Ghosh, S. S., Goulas, A., Falkiewicz, M., Huntenburg, J. M., Langs, G., Bezgin, G., Eickhoff, S. B., Castellanos, F. X., Petrides, M., et al. (2016). Situating the default-mode network along a principal gradient of macroscale cortical organization. Proceedings of the National Academy of Sciences, 113 (44), 12574–12579.

Medaglia, J. D., Huang, W., Karuza, E. A., Kelkar, A., Thompson-Schill, S. L., Ribeiro, A., & Bassett, D. S. (2018). Functional alignment with anatomical networks is associated with cognitive flexibility. Nature human behaviour, 2 (2), 156–164.

Medaglia, J. D., Lynall, M.-E., & Bassett, D. S. (2015). Cognitive network neuroscience. Journal of cognitive neuroscience, 27 (8), 1471–1491.

Mediano, P. A., Rosas, F. E., Timmermann, C., Roseman, L., Nutt, D. J., Feilding, A., Kaelen, M., Kringelbach, M. L., Barrett, A. B., Seth, A. K., et al. (2024). Effects of external stimulation on psychedelic state neurodynamics. ACS Chemical Neuroscience, 15 (3), 462–471.

Meer, J. N. v. d., Breakspear, M., Chang, L. J., Sonkusare, S., & Cocchi, L. (2020). Movie viewing elicits rich and reliable brain state dynamics. Nature communications, 11 (1), 5004.

Mesulam, M.-M. (1998). From sensation to cognition. Brain: a journal of neurology, 121 (6), 1013–1052.

Nentwich, M., Ai, L., Madsen, J., Telesford, Q. K., Haufe, S., Milham, M. P., & Parra, L. C. (2020). Functional connectivity of eeg is subject-specific, associated with phenotype, and different from fmri. NeuroImage, 218, 117001.

Pang, J. C., Aquino, K. M., Oldehinkel, M., Robinson, P. A., Fulcher, B. D., Breakspear, M., & Fornito, A. (2023a). Geometric constraints on human brain function. Nature, 1–9.

Pang, J. C., Aquino, K. M., Oldehinkel, M., Robinson, P. A., Fulcher, B. D., Breakspear, M., & Fornito, A. (2023b). Reply to: Commentary on pang et al.(2023) nature. bioRxiv, 2023–10.

Park, H.-J., & Friston, K. (2013). Structural and functional brain networks: From connections to cognition. Science, 342 (6158), 1238411.

Pascual-Marqui, R. D., Lehmann, D., Koukkou, M., Kochi, K., Anderer, P., Saletu, B., Tanaka, H., Hirata, K., John, E. R., Prichep, L., et al. (2011). Assessing interactions in the brain with exact low-resolution electromagnetic tomography. Philosophical Transactions of the Royal Society A: Mathematical, Physical and Engineering Sciences, 369 (1952), 3768–3784.

Pascucci, D., Rubega, M., Rué-Queralt, J., Tourbier, S., Hagmann, P., & Plomp, G. (2022). Structure supports function: Informing directed and dynamic functional connectivity with anatomical priors. Network Neuroscience, 6 (2), 401–419.

Pascucci, D., Tourbier, S., Rué-Queralt, J., Carboni, M., Hagmann, P., & Plomp, G. (2022). Source imaging of high-density visual evoked potentials with multi-scale brain parcellations and connectomes. Scientific Data, 9 (1), 9.

Patil, K. R., Jung, K., & Eickhoff, S. B. (2023). Commentary on pang et al.(2023) nature. bioRxiv, 2023–10.

Pirondini, E., Vybornova, A., Coscia, M., & Van De Ville, D. (2016). A spectral method for generating surrogate graph signals. IEEE signal processing letters, 23 (9), 1275–1278.

Preti, M. G., & Van De Ville, D. (2019). Decoupling of brain function from structure reveals regional behavioral specialization in humans. Nature communications, 10 (1), 4747.

Rigoni, I., Queralt, J. R., Glomb, K., Preti, M., Roehri, N., Tourbier, S., Spinelli, L., Seeck, M., Van De Ville, D., Hagmann, P., et al. (2023). Structure-function coupling increases during interictal spikes in temporal lobe epilepsy: A graph signal processing study. Clinical Neurophysiology.

Rué-Queralt, J., Glomb, K., Pascucci, D., Tourbier, S., Carboni, M., Vulliémoz, S., Plomp, G., & Hagmann, P. (2021). The connectome spectrum as a canonical basis for a sparse representation of fast brain activity. NeuroImage, 244, 118611.

Rué-Queralt, J., Mancini, V., Rochas, V., Latrèche, C., Uhlhaas, P. J., Michel, C. M., Plomp, G., Eliez, S., & Hagmann, P. (2023). The coupling between the spatial and temporal scales of neural processes revealed by a joint time-vertex connectome spectral analysis. NeuroImage, 280, 120337.

Salo, T., Yarkoni, T., Nichols, T. E., Poline, J.-B., Kent, J. D., Gorgolewski, K. J., Glerean, E., Bottenhorn, K. L., Bilgel, M., Wright, J., Reeders, P., Kimbler, A., Nielson, D. N., Yanes, J. A., Pérez, A., Oudyk, K. M., Jarecka, D., Enge, A., Peraza, J. A., & Laird, A. R. (2022, January). Neurostuff/nimare: 0.0.11 (Version 0.0.11). Zenodo. 10.5281/zenodo.5826281

Samara, A., Eilbott, J., Margulies, D. S., Xu, T., & Vanderwal, T. (2023). Cortical gradients during naturalistic processing are hierarchical and modality-specific. NeuroImage, 271, 120023.

Shuman, D. I., Narang, S. K., Frossard, P., Ortega, A., & Vandergheynst, P. (2013). The emerging field of signal processing on graphs: Extending high-dimensional data analysis to networks and other irregular domains. IEEE signal processing magazine, 30 (3), 83–98.

Smith, R. E., Tournier, J.-D., Calamante, F., & Connelly, A. (2015). Sift2: Enabling dense quantitative assessment of brain white matter connectivity using streamlines tractography. Neuroimage, 119, 338–351.

Spielman, D. A. (2007). Spectral graph theory and its applications. 48th Annual IEEE Symposium on Foundations of Computer Science (FOCS’07), 29–38.

Tewarie, P., Abeysuriya, R., Byrne, Á., O’Neill, G. C., Sotiropoulos, S. N., Brookes, M. J., & Coombes, S. (2019). How do spatially distinct frequency specific meg networks emerge from one underlying structural connectome? the role of the structural eigenmodes. NeuroImage, 186, 211–220.

Tournier, J.-D., Smith, R., Raffelt, D., Tabbara, R., Dhollander, T., Pietsch, M., Christiaens, D., Jeurissen, B., Yeh, C.-H., & Connelly, A. (2019). Mrtrix3: A fast, flexible and open software framework for medical image processing and visualisation. Neuroimage, 202, 116137.

Xu, H., Lorbert, A., Ramadge, P. J., Guntupalli, J. S., & Haxby, J. V. (2012). Regularized hyperalignment of multi-set fmri data. 2012 IEEE statistical signal processing workshop (SSP), 229–232.

Yang, Y., Zheng, Z., Liu, L., Zheng, H., Zhen, Y., Zheng, Y., Wang, X., & Tang, S. (2023). Enhanced brain structure-function tethering in transmodal cortex revealed by high-frequency eigenmodes. Nature Communications, 14 (1), 6744.

Yarkoni, T., Poldrack, R. A., Nichols, T. E., Van Essen, D. C., & Wager, T. D. (2011). Large-scale automated synthesis of human functional neuroimaging data. Nature methods, 8 (8), 665–670.

Yeo, B. T., Krienen, F. M., Sepulcre, J., Sabuncu, M. R., Lashkari, D., Hollinshead, M., Roffman, J. L., Smoller, J. W., Zöllei, L., Polimeni, J. R., et al. (2011). The organization of the human cerebral cortex estimated by intrinsic functional connectivity. Journal of neurophysiology.

Zamora-López, G., & Gilson, M. (2024). An integrative dynamical perspective for graph theory and the analysis of complex networks. Chaos: An Interdisciplinary Journal of Nonlinear Science, 34 (4).

Zhou, D., Zhang, G., Dang, J., Unoki, M., & Liu, X. (2022). Detection of brain network communities during natural speech comprehension from functionally aligned eeg sources. Frontiers in Computational Neuroscience, 16, 919215.

